# Adaptation of the *Chlorella* photosynthetic electron transport chain to environmental light conditions

**DOI:** 10.1101/2025.05.13.653806

**Authors:** Grant Steiner, Devinjeet Saini, Arivu Kapoor, Colin Gates

## Abstract

In photosynthetic organisms, the process of photosystem II-cyclic electron flow (PSII-CEF) protects against degradation of the active D1 protein subunit of photosystem II in high light environments. By comparing the photophysiology of the extreme-light (2000 µEin/m^2^/s) desert alga *Chlorella ohadii* to the low-light (20 µEin/m^2^/s) aquatic alga *Chlorella* sp. NIES 642, we can monitor how natural adaptations resulting in differing levels of PSII-CEF affect the photosynthetic electron transport chain. Modeling of chlorophyll fast repetition rate fluorometry shows distinct backward transitions of the Kok cycle in *C. ohadii*, which are absent in NIES 642. Increase in PSII-CEF also has a positive effect on the occurrence of miss parameters in the water oxidizing complex. Fluorescent Q _A_^-^ reoxidation kinetics detail that the majority of reaction centers in *C. ohadii* are performing electron transfer to oxidized Q_B_ under saturating light conditions. A combination of kinetic observations excludes plastocyanin as a potential external electron carrier in the mechanistic path of PSII-CEF and suggests that photosystem I-cyclic electron flow works in tandem with PSII-CEF. The combination of these two alternative flow mechanisms expedite electron transfer downstream of PSII and optimize ATP production, respectively. Utilizing 77K spectrofluorometry, congruent photosystem stoichiometry is found between both *Chlorella* species, despite a 100-fold growth light intensity difference. Electrochromic shift measurements show that *C. ohadii* has diminished changes to both trans-membrane potential and ΔpH during operation of photosynthesis compared to NIES 642, and that excess addition of N,N’-dicyclohexylcarbodiimide to *Chlorella* cells has an inhibitory effect on the photosynthetic electron transport chain.

## 1. Introduction

For many organisms capable of performing oxygenic photosynthesis, one hindrance to their continued survival is the inability to relocate or shelter themselves in response to environmental stressors, frequently including light and drought stress[1]. To counteract this lack of mobility, plants, algae, and cyanobacteria have independently evolved various biochemical adaptations to their photosynthetic electron transport chain (PETC)[2-4]. Preserving the integrity of the PETC is of utmost importance to phototrophic organisms as this machinery acts as a biological transducer to derive chemical energy, namely NADPH and ATP, from light energy. Various protective mechanisms of the PETC are moderately understood that employ alternative pathways of electron transport differing from the typical linear flow from water to NADPH. In cyclic electron flow around photosystem I (PSI), the creation of NADPH is circumvented without sacrificing production of ATP[5, 6]. In various organisms, this is accomplished by the *proton gradient regulation 5* protein returning electrons to cytochrome (cyt) *b*_*6*_*f* from ferredoxin[7, 8]. In the mechanism of pseudo-cyclic electron flow, or the water-water cycle, electrons leave the reaction center of PSI and are used by a family of enzymes known as flavodiiron proteins to reduce molecular oxygen to water, the inverse of the primary step of linear electron flow[9]. Plastoquinol terminal oxidase pathways also consume oxygen to create water, directly utilizing electrons from reduced plastoquinone (PQ) within the thylakoid membrane[10, 11]. One valuable photoprotective mechanism involves redirecting the flow of electrons to cycle around the primary major photosynthetic protein in the PETC, photosystem II (PSII), when excess photons are present[12]. This process of PSII-cyclic electron flow (PSII-CEF) protects the active D1 protein subunit of PSII and ultimately the phototroph itself from the production of singlet oxygen and excess high energy reactive oxygen species[13]. Unlike the other methods of alternative electron transport in the PETC previously mentioned, the mechanism of action for PSII-CEF is not fundamentally understood[14].

What is known to date about PSII-CEF is that charge-separated electrons are delivered to the plastoquinone binding site Q_B_, the site of terminal electron acceptance within PSII, located near the stromal surface of the D1 subunit; electrons are then returned to the donor side of PSII, the water oxidizing complex (WOC)[15]. The acceptor side of PSII these recycled electrons originate from comprises an irreversibly bound plastoquinone (Q_A_) that sequentially transfers one electron at a time past a non-heme iron to a reversibly bound plastoquinone (Q_B_). Q_B_ accepts two electrons, balanced by acceptance of two protons from the stromal side of the thylakoid, forming plastoquinol (PQH_2_)[16]. PQH_2_ carries reducing potential from PSII to the inter-membrane space of the thylakoid and to the cyt *b*_*6*_*f* complex, where one electron is sent linearly through cyt *f*, and one re-reduces PQ in a quinone cycle (Q-cycle) via transfer through cyt *b*_*6*_[17]. In the cyanobacterium *Thermosynechococcus vulcanus*, a putative third PQ binding site, Q_C_, was observed lumenal to Q_B_, with an approximate distance of 12 Å between the quinone head groups[18]. The appearance and prevalence of the Q_C_ site seems to be dependent on preparation methods[19] possibly due in part to the lower binding affinity of the polar quinone head group than in the Q_B_ site[20]. Q_C_ has been theorized to simply modulate high and low potential redox forms of cyt *b*_*559*_[21], but it shows distinct advantages as a potential electron carrier within an internal mechanism of PSII-CEF. PSII-CEF bypasses the use of water for electrons, and possibly for protons as well, as it has been posited to be a proton-coupled electron transfer (PCET) process that optimizes ATP production by increasing the trans-thylakoid proton gradient[22]. Recent studies of flash-induced oximetry on *T. elongatus* microcrystals suggest a switch from linear electron flow to PSII-CEF is dependent on occupancy of the Q_C_ site, which would regulate electron allocation between Q_C_-mediated single-electron return to the WOC, and the typical two-electron transfer to PQH_2_, which likely also returns electrons to the WOC with an accompanying two-proton deposition into the lumen[23]. These hypothesized mechanisms, illustrated in **Fig. 1**, all include electron return to the WOC.

**Figure 1.**
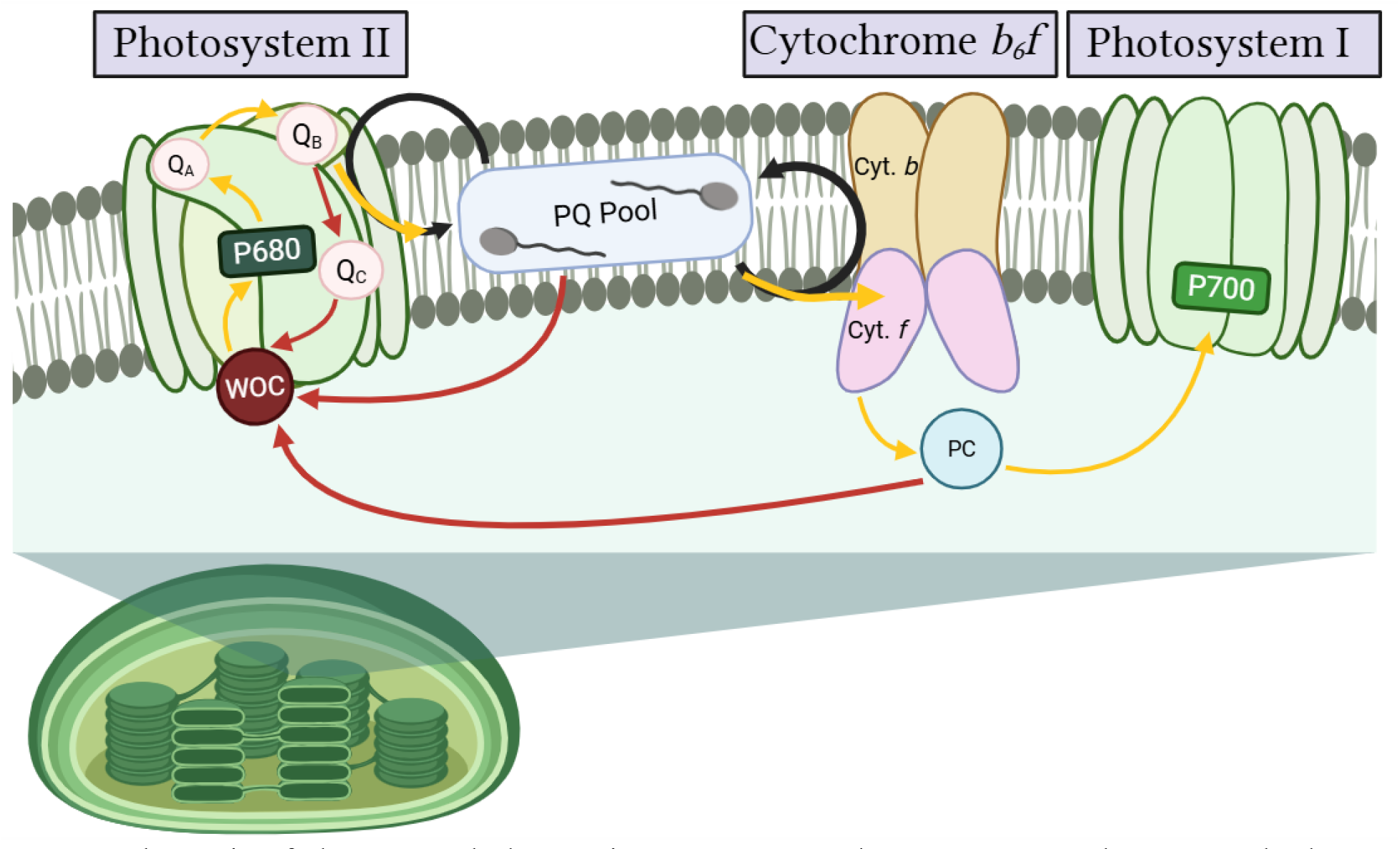
Schematic of electron and plastoquinone movement between PSII and PSI. Standard linear electron transfer is shown with yellow arrows between intermediates. Plastoquinone movement within the thylakoid membrane is shown in black arrows, and three possible PSII-CEF pathways are shown in red arrows. These potential pathways include an internal mechanism through the putative quinone binding site Q_C_, directly from reduced forms of PQ, or through the lumenal surface of the thylakoid membrane via transfer from plastocyanin (PC).

These potential terminal electron donors in PSII-CEF possess both spatial access and sufficient redox potential to reduce the WOC. The WOC, the primary PSII electron acceptor, is a heterocubane calcium-manganese cluster with five oxo bridges, complexed to four water ligands and the protein scaffold (Formula: Mn_4_CaO_5_)[19]. Water arrives at the WOC through nearby water channels[24, 25], and is split into electrons, protons, and molecular oxygen at the catalytic site. WOC efficiency ensures electron input to the PETC, although it is generally not the rate-limiting step at the photosynthetic optimum[26, 27]. Up to three oxidizing equivalents are semi-stably retained in the WOC by oxidizing the manganese atoms, producing a cycle (the Kok cycle) of WOC intermediates known as S-states (oxidation states)[28]. There are five S-states of the WOC, S_0_-S_4_, where the subscript denotes the number of electrons removed from the WOC[29, 30]. S_0_ and S_1_ are the only dark-stable WOC intermediates, as S_2_ and S_3_ are unstable over minute time scales[31, 32] and decay back to S_1_, while S_4_ decays forward to S_0_ faster than observable by extant methods, including femtosecond X-ray crystallography[33]. When the S_4_ state decays, it releases molecular oxygen, coordinates to water, and returns to S_0_[34]. One advantage to observation of the PSII donor side is that the S-states of the WOC vary in their reaction center fluorescence, due to shifting midpoint potential gaps, causing corresponding fluctuations in chlorophyll *a* variable fluorescence intensity[35, 36]. Kok cycle-derived matrix models, such as the VZAD model used herein, analyze these oscillations in PSII variable fluorescence data and determine amplitudes of WOC cycle efficiency[37]. VZAD calculates parameters of WOC inefficiency that describe scenarios other than the advance of a single electron from the WOC. Alpha is a measure of failure to net advance an electron from the WOC (miss), beta measures advancing two electrons from the WOC (double hit), and delta measures electron return to either the S_2_ or S_3_ states of the WOC (backward transition). Delta is a direct measure of PSII-CEF, as it is a consequence of electron return to the WOC[15]. The modified Kok cycle and attributed inefficiencies monitored by the VZAD model are illustrated in **Supplemental Figure S1**.

To experiment on systems that are genetically implementable in higher order plants, namely field crops, we have initiated our mechanistic inquiry into PSII-CEF utilizing green algal strains. Green algae are most closely related to plants in terms of cell structure and mechanism of photosynthetic function, but can proliferate exponentially faster than plants[38]. *Chlorella* is a model algal genus that has shown reported variance in growth and PSII-CEF in response to light[32, 39, 40]. Many strains in this genus are of biotechnological and/or scientific interest. For comparison of environmental adaptation, two *Chlorella* strains were grown and tested. *Chlorella ohadii* is an arid, high-light adapted strain that possesses the fastest known doubling time of any phototrophic organism at 1.4 hours and shows high levels of PSII-CEF, around 90%[32]. *C. ohadii* grows in desert soil crusts (Negev, Israel) that receive extreme amounts of solar radiation, creating the environmental necessity of photo-protecting PSII[22]. The environment in which *C. ohadii* natively grows exhibits conditions of highly varying temperature, salinity, and hydration, and this organism performs unprecedented levels of photoprotection without elevating levels of non-photochemical quenching[41]. *C. ohadii* possesses novel photoprotective proteins[42] and performs varying levels of PSII-CEF dependent on light intensity, peaking at its native environmental conditions (2000 µEin/m^2^/s)[32]. Conversely, its relative *Chlorella* sp. NIES 642 (hereafter NIES 642) is an aquatic, low-light adapted strain, native to the Miyata River in Ibaraki, Japan[43]. NIES 642 has approximately 1% the environmental light intensity (20 µEin/m^2^/s) compared to *C. ohadii*, as it is shaded by natural foliage, and experiences far less extreme thermal fluctuations. Due to the exponentially lower solar radiation, NIES 642 has a much larger light harvesting complex for PSII[44] and reduced needs of photoprotection compared to *C. ohadii*[40]. These environmental and physiological distinctions make NIES 642 an advantageous model of low PSII-CEF for comparison.

## 2. Materials and Methods

### 2.1 Culture growth

All cultures were grown within a CARON Model 7314-22 plant growth chamber set to a temperature of 30°C. Atmospheric carbon dioxide levels within the chamber were regulated to 1.5%. Algal cultures were grown in 100 mL of BG-11 medium at pH 7.5[45, 46] within 250 mL Erlenmeyer flasks. Both *Chlorella* species were grown at their natural light intensities. These intensities, determined by a LI-COR Biosciences LI-250A light meter, correspond to a photon delivery of 20 µEin/m^2^/s (NIES 642)[43] and 2000 µEin/m^2^/s (*C. ohadii*)[40]. Logarithmic growth times were determined by rate of change analysis on measurements of optical density at 730 nm (OD_730_) using a Thermo-Fisher Genesys 10 spectrophotometer. All culture testing was performed during log-phase growth and normalized to equivalent OD_730_ between strains for all testing methods. Comparative growth curves for both *Chlorella* species are reported in **Supplemental Figure S2**.

### 2.2 Fast repetition rate (FRR) fluorometry

Cultures were dark-adapted for 180s, allowing the WOCs in each PSII to decay to the dark-stable S-states, S_0_ and S_1_[34]. Utilizing a modified Joliot-type spectrometer (JTS-150) from SpectroLogiX equipped with a 5-watt fiber output laser (RPMC Lasers) at a central wavelength of 636 nm and spectral width of 1.1 nm, single turnover flashes (STFs) were created via a 24 μs laser pulse to saturate the culture and advance all PSII centers by one S-state[35, 47]. During the saturating flash, rising fluorescence intensity was detected every 0.6 µs. All FRR measurements were performed at a frequency of 10 Hz (100 ms between STFs) to detect changes in variable fluorescence without giving the WOC time to decay in higher oxidation states.

Background fluorescence of the system (F_o_) was subtracted from the maximal fluorescence reached (F_m_) to acquire the variable fluorescence, F_v_, for each STF[48]. A group of 50 STFs (flash train) was recorded and replicated 14 more times, with a 120s wait time between each flash train to re-induce the dark stable population distribution[36]. Data recorded from the JTS-150 was processed by MATLAB r2023a software to produce F_v_/F_m_ ratios for each STF. These data were fitted by the model-dependent nonlinear least-squares matrix model VZAD[37], as average F_v_/F_m_ values for the 15 flash trains. The VZAD package used to average inefficiency parameters of the WOC in this work was version 4.0, run using PyCharm 2024.2.1. FRR testing cultures were normalized to an OD_730_ of 0.30.

### 2.3 Rate oximetry by chlorophyll concentration

Oxygen evolution rates were obtained using a Hansatech Oxygraph+ Clark-type electrode containing a central platinum cathode encircled by a silver anode, bridged with 50% KCl and covered with a paper spacer/polytetrafluoroethylene membrane[49]. Respiration was measured under darkness for five minutes before replacement with fresh sample and oxygen evolution measurements taken under illumination equal to the culture’s growth light intensity. Rate of change for both respiration and oxygen evolution were determined by linear fit analysis in Origin 2023b. Overall oxygen evolution was calculated by difference of respiratory rate from oxygen evolving rate.

Rate oximetry values were quantified in μmol O_2_/mg chl *a*/h via chlorophyll extraction. Chlorophyll *a* concentrations were determined according to the extinction coefficients for chlorophyll extraction in chilled methanol established by Porra et. al (1989)[50]. Chlorophyll measurements were performed in both biological and technical triplicate (n=9). Rate oximetry and chlorophyll extraction measurements were both normalized to an OD_730_ of 0.60.

### 2.4 Q_A_^-^ reoxidation kinetics

The modified JTS-150 was also utilized to monitor PSII acceptor side kinetics via chlorophyll fluorescence changes by the two-period behavior of electron transfer to plastoquinone. Following protocol of Gorbunov et al. (1999)[47] an over-saturating flash of 58.5 µs duration provided an overall F_o_ and F_m_ of the culture, to which decaying background fluorescence over time (F_n_) was compared. Following the saturating flash was a 5.4 µs dark period before a second STF, from which a new background fluorescence (F_1_) was recorded. The dark period until the second STF was increased to 8.4 µs to record F_2_, to 12.4 µs to record F_3_, and increased repeatedly until a final dark time of 8.19 ms (F_14_). Individual F_v_/F_m_ values for each STF were plotted against their respective dark period length in μs, using an average of triplicate datasets. These variable fluorescence changes were fit to a biphasic decay regression analysis in Origin 2023b, from which kinetic parameters of Q_A_^-^ oxidation amplitude were extrapolated. Q_A_^-^ reoxidation testing cultures were normalized to an OD_730_ of 0.10 to ensure complete initial reduction of plastoquinone at the Q_A_ binding site.

### 2.5 Cytochrome *b*_*6*_*f* and plastocyanin redox kinetics

The absorption of reduced forms of plastocyanin, cyt *b*_*6*_, and cyt *f* were measured during an illumination window with a duration of five seconds. The modified JTS-150 supplied actinic light at 630 nm and an intensity of 3200 µEin/m^2^/s to energetically saturate the electron transport chain, with margins of dark time before and after the illumination window for baseline measurements and re-equilibration observations, respectively. These dark times enclosing the illumination window are conserved for all absorbance measurements on the JTS-150. During the light interval, multiple single wavelength LEDs were monitored to observe the redox kinetics of the copper center of PC and the heme centers in cyt *b*_*6*_ and cyt *f*. These PETC intermediates were simultaneously measured at 574 nm, 563 nm, and 554 nm, respectively, with a baseline corrective measurement at 546 nm. Coefficients for correction of spectral overlap of artifacts, integrated by the PhotoKine software utilized by the JTS-150, are available in the associated SpectroLogiX user manual for the instrument. A multiple bandpass filter (BG-39) was placed within the beam path preceding 1 mL of sample within a semi-micro cuvette with clear windows parallel to the beam path. To ensure baseline corrected measurements, all LEDs in use were balanced to an output voltage of 6.5 V. All absorbance testing cultures herein (cyt *b*_*6*_*f*, ECS, and P700) were normalized to an OD_730_ of 1.00, with results reported being representative of a triplicate of technical trials.

### 2.6 77K spectrofluorometry

Fluorescence emission spectra were taken using a JASCO FP-8300 spectrofluorometer with a specialized liquid nitrogen dewar for cryostatic samples. Cultures were placed in 7” medium wall glass NMR tubes and flash frozen before placement into the spectrofluorometer. Chlorophyll *a* fluorescence was monitored with an excitation wavelength of 435 nm[51]. The resolution speed of the spectrofluorometer was set to 500 nm/min. All 77K testing cultures were normalized to an OD_730_ of 0.60, with a single test being representative of a triplicate of technical trials. Gaussian deconvolution of the 77K spectra was performed using the Peak Pick analysis function within OriginLab 2023b to obtain relative abundance of chlorophyll by photosystem association within both *Chlorella* species.

### 2.7 P700 absorbance

Absorbance changes of the P700 special chlorophyll *a* pair in PSI can be monitored in the near-infrared region of the electromagnetic spectrum[52]. Unlike previously discussed absorbance studies on the JTS-150, P700 measurements were run with a different multiple bandpass filter preceding the detector and single wavelength lamps of 810 nm to monitor PSI band-shift, with control measurements at 705 and 740 nm to reduce interference from measurements of plastocyanin.

### 2.8 Electrochromic shift (ECS)

Electrochromic shift (ECS) was utilized as a means of monitoring trans-membrane electrochemical potential and proton gradient by observation of absorption changes in thylakoid membrane pigments with and without an external light source[53]. ECS testing uses the same hardware and timescale as cyt *b*_*6*_*f* measurements on the JTS-150 (actinic light of 630 nm at an intensity of 3200 µEin/m^2^/s in 5 second dark-light-dark intervals with a BG-39 bandpass filter). For ECS, the wavelengths that were monitored have been determined by diffused-optics flash spectroscopy (DOFS)[54]. These wavelengths are 520 nm to monitor ECS, and 546 nm to monitor the light scattering properties of the thylakoid membrane. DOFS shows that 520 nm in plants and green algae is a combined contribution of 100% ECS signal and 72% scattering signal, while 546 nm is a combination of 93% scattering signal and 49% ECS signal. These two wavelengths were again balanced to an output voltage of 6.5 V, and the DOFS attained coefficients were processed by the JTS-150 to produce traces of ECS. N,N’-dicyclohexylcarbodiimide (DCCD) testing was performed by adding 100 μM DCCD, well above the known inhibitory concentration of *Chlorella* cells[55], into 1.00 OD_730_ culture in the absence of light.

## 3. Results

### 3.1 Fast repetition rate fluorometry (FRR)

Oscillatory behavior of variable chlorophyll fluorescence within the PSII reaction centers of *C. ohadii* and NIES 642 was promoted by a series of STFs, and is reported in **Fig 2a + b**, respectively. An important metric retrieved from FRR testing is the WOC quality factor (Q), the reciprocal of the sum of all WOC progression inefficiencies, all of which are reported in **Table 1**. The reported Q values, 4.33 in the high PSII-CEF model, and 6.80 in the low PSII-CEF, approximate the amount of quality period-four oscillations that occur prior to fluorescence quenching, i.e. the “quality” of S-state advancement and retention of population distribution. Visibly, this can be observed by the fluorescence of *C. ohadii* reaction centers equilibrating after approximately 17 STFs, and NIES 642 equilibrating after approximately 27 STFs. The lower quality of oscillatory behavior in *C. ohadii* stems from the higher occurrence of misses (alpha), double hits (beta), and backward transitions (delta) in S-states. Backward transitions are an integral consequence of PSII-CEF, as they describe direct electron return to the WOC during the S_2_ and S_3_ states, resulting in the reduction of the WOC[15, 32, 37, 56]. Computationally, backward transitions are absent in the low PSII-CEF model, and most of the inefficient transitions associated with NIES 642 stem from misses. Misses are also elevated in *C. ohadii*; however, this increase in failures to advance may be artificially high, due to successful electron advance in combination with an electron return to the donor side of PSII, resulting in the appearance of an unchanged variable chlorophyll fluorescence, while masking a further increase in the delta parameter. Further deconvolution with a modified model may be required to separate these two instances.

**Table 1.**
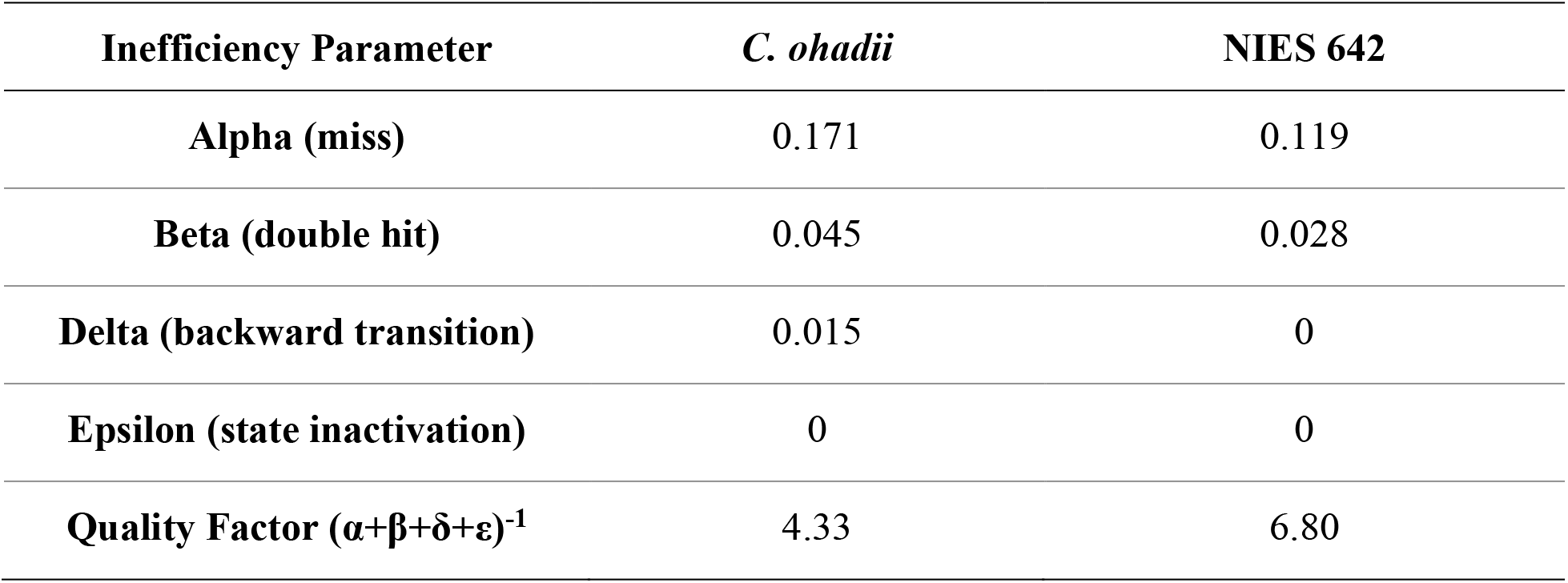
WOC inefficiency parameters calculated from variable chlorophyll *a* fluorescence oscillations detailed in Fig. 2.

**Figure 2.**
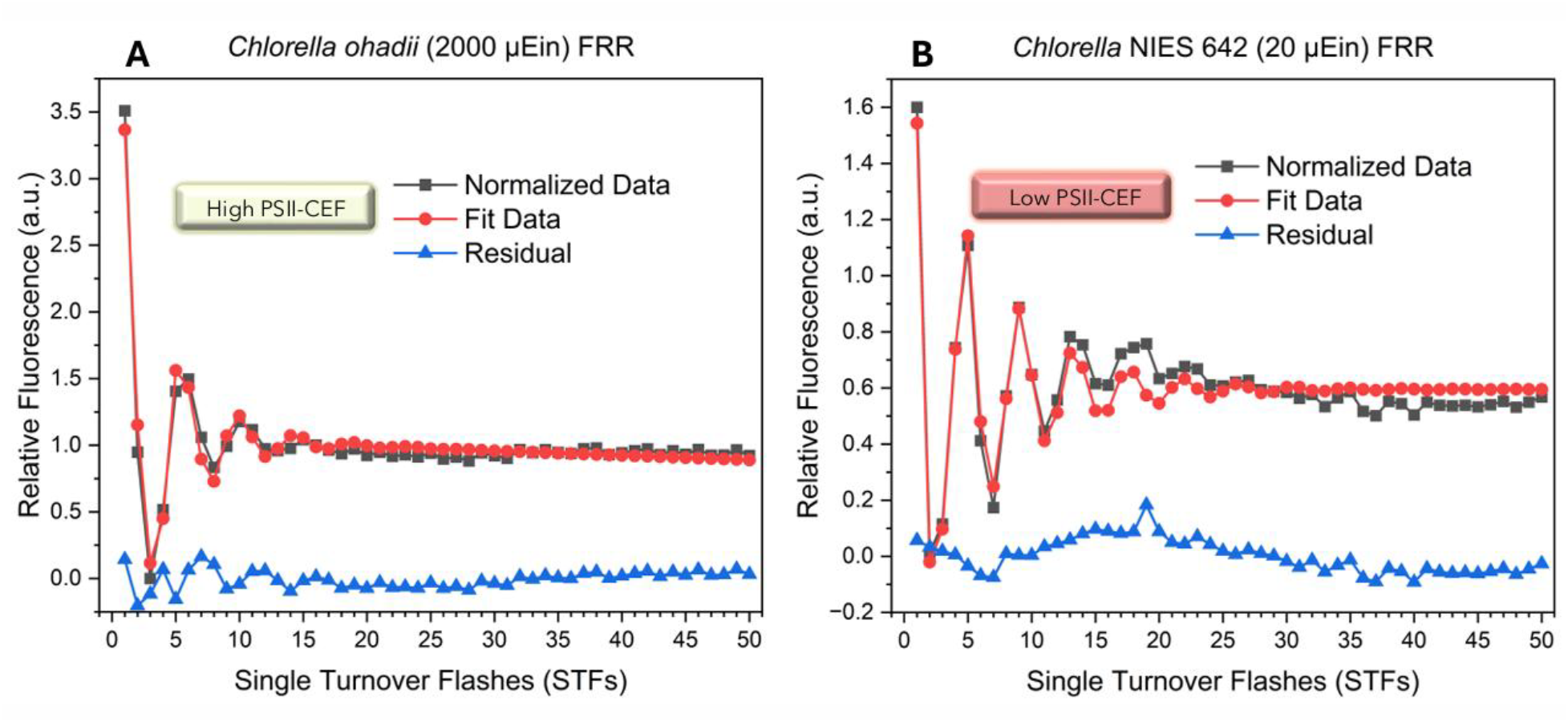
FRR variable chlorophyll *a* fluorescence (F_v_/F_m_) ratios normalized (black trace) and fit (red trace) by the VZAD software for both *Chlorella* strains. The residual (blue trace) is utilized to produce quantities of inefficient transitions for **a)** *C. ohadii*, the model of high PSII-CEF, and **b)** NIES 642, the low PSII-CEF model.

The diminished WOC efficiency in the high PSII-CEF model is to be expected, as an overabundance of excitons being created by the PSII reaction center cannot be removed continuously. In turn, without the creation of the electron hole in P680, the oxidizing equivalents being stored in the WOC are not removed as effectively as they are in the low PSII-CEF model. These contrasting WOC efficiencies are further validated by experimental rate oximetry testing. In environmental light intensities, *C. ohadii* exhibits an average oxygen evolution rate of 1254 μmol O_2_/mg chl *a*/h, and NIES 642 evolves 41.4 μmol O_2_/mg chl *a*/h. Considering the 1% relative photon incidence in the low PSII-CEF conditions, NIES 642 has approximately 3.3-fold higher per-chlorophyll quantum efficiency vs. *C. ohadii*, consistent with the relative comparisons of WOC inefficiencies found in FRR and the previously observed behavior of *C. ohadii* at varying light intensities[32]. Complete oxygen rates and chlorophyll extraction data are available in **Supplementary Figure S3**.

### 3.2 Q_A_^-^ reoxidation kinetics

A running set of STFs at increasing time intervals between them can monitor the two-period behavior of electron transfer from the plastoquinone Q_A_ in whole algal and plant cells[47, 57, 58]. An initial, saturating light pulse is used to fully reduce the Q_A_ site. Continuing with a series of pulses can quantify the kinetics of Q_A_^-^ reoxidation while avoiding actinic behavior[47] and non-linear effects of photosystem-antenna connections so that both *Chlorella* may be contrasted, despite the stark difference in antenna size at their environmental light conditions. Regression analysis of the proceeding decay in reaction center fluorescence by changes in acceptor side redox potential (**Fig. 3**) quantifies kinetic parameters of plastoquinone electron transfer (**Table 2**). These kinetic parameters are invaluable towards elucidating the origin of the electrons returned to the WOC, as most literature information to date regarding the potential mechanisms of PSII-CEF involve some form of modulation of electron transfer through either Q_A_ or Q_B_[23, 32, 59-61].

**Table 2.** Parameters of PSII acceptor side electron transfer kinetics, retrieved from analysis of the decay traces in Fig. 3.

| Inefficiency Parameter | *C. ohadii* | NIES 642 |
| --- | --- | --- |
| **y<sub>0</sub>** | 7.6% | 14.9% |
| **t<sub>1</sub>** | 77 μs | 105 μs |
| **A<sub>1</sub>** | 53.7% | 21.8% |
| **t<sub>2</sub>** | 1.25 ms | 2.20 ms |
| **A<sub>2</sub>** | 38.7% | 63.3% |

**Figure 3.**
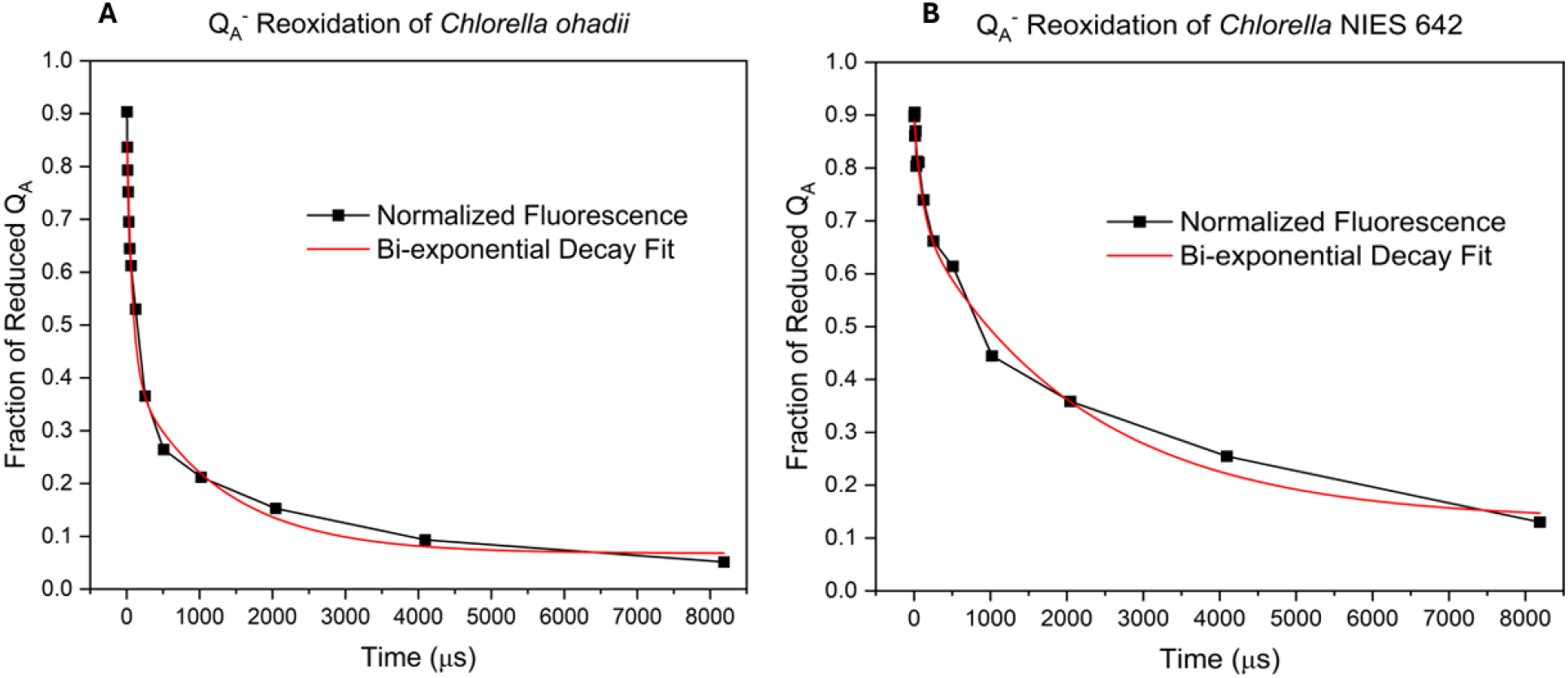
Biphasic decay fitting of Q_A_^-^ population in reaction centers of **a)** *C. ohadii* and **b)** NIES 642.

Comparison of the two PSII-CEF models illustrates stark differences in the redox poise of their plastoquinone pools. t_1_ reflects the time for Q_A_^-^ to reduce Q_B_, while A_1_ is the fraction of reaction centers with Q_B_ attached. t_2_ and A_2_ are analogous to t_1_ and A_1_, with the replacement of Q_B_ for semiquinone (Q_B_•-). y_0_ reflects the remaining centers, with either no bound quinone or bound PQH_2_. This population of centers is dependent on the rate limitation of Q_B_ replacement, the rate limiting step of PSII electron transfer, which has an approximate diffusion time of 3-5 ms[62-64]. In *C. ohadii*, we observe approximately half the quantity of unbound quinone population (7.6%) compared to NIES 642 (15%, which is itself relatively low[65-67]), showing almost complete utilization of the PQ pool in both species of *Chlorella*.

Reoxidation of Q_A_^-^ occurs within a few hundred microseconds in a majority of *C. ohadii* reaction centers, while both *Chlorella* species have comparable, atypically quick, reduction of oxidized Q_B_. Despite this similarity in a small t_1_, the species have a notably disparate fraction of centers performing the primary electron transfer. The largest inter-species discrepancy occurs around 500 μs of illumination, where only 26% of centers in *C. ohadii* possess reduced Q_A_, while this quantity is 61% in NIES 642. *C. ohadii* also exhibits a smaller population and shorter lifetime of the semiquinone form available in the Q_B_ site (38.7%). This is not the case for NIES 642, where a majority of the PSII centers (63.3%) exhibit stable, long-lived Q_B_•^-^ binding. These differences culminate in a significantly higher utilization of the PQ pool in the high PSII-CEF system.

### 3.3 Cytochrome *b*_*6*_*f* absorbance

As the redox bridge between the photosystems and the major protein that immediately follows PSII in linear electron flow, it is advantageous to monitor the redox kinetics of the cyt *b*_*6*_*f* protein to judge electron input following PSII itself. The dimeric cyt *b*_*6*_*f* protein contains 4 prosthetic heme groups and an iron-sulfur cluster in each monomer that ultimately shuttle electrons from the product of PSII redox reactions, PQH_2_, to another free electron carrier, PC[68]. Investigation of PC in particular is necessitated by its redox potential[69] and spatial availability of electron donation to the WOC via diffusion to the lumenal surface of PSII. Absorbance spectra monitoring the change in redox forms of PC (oxidized form - reduced form), along with cyt *b*_*6*_ and cyt *f* (reduced form – oxidized form) can directly contrast cyt *b*_*6*_*f* electron transfer amplitudes between the models of PSII-CEF and are reported in **Fig. 4**.

**Figure 4.**
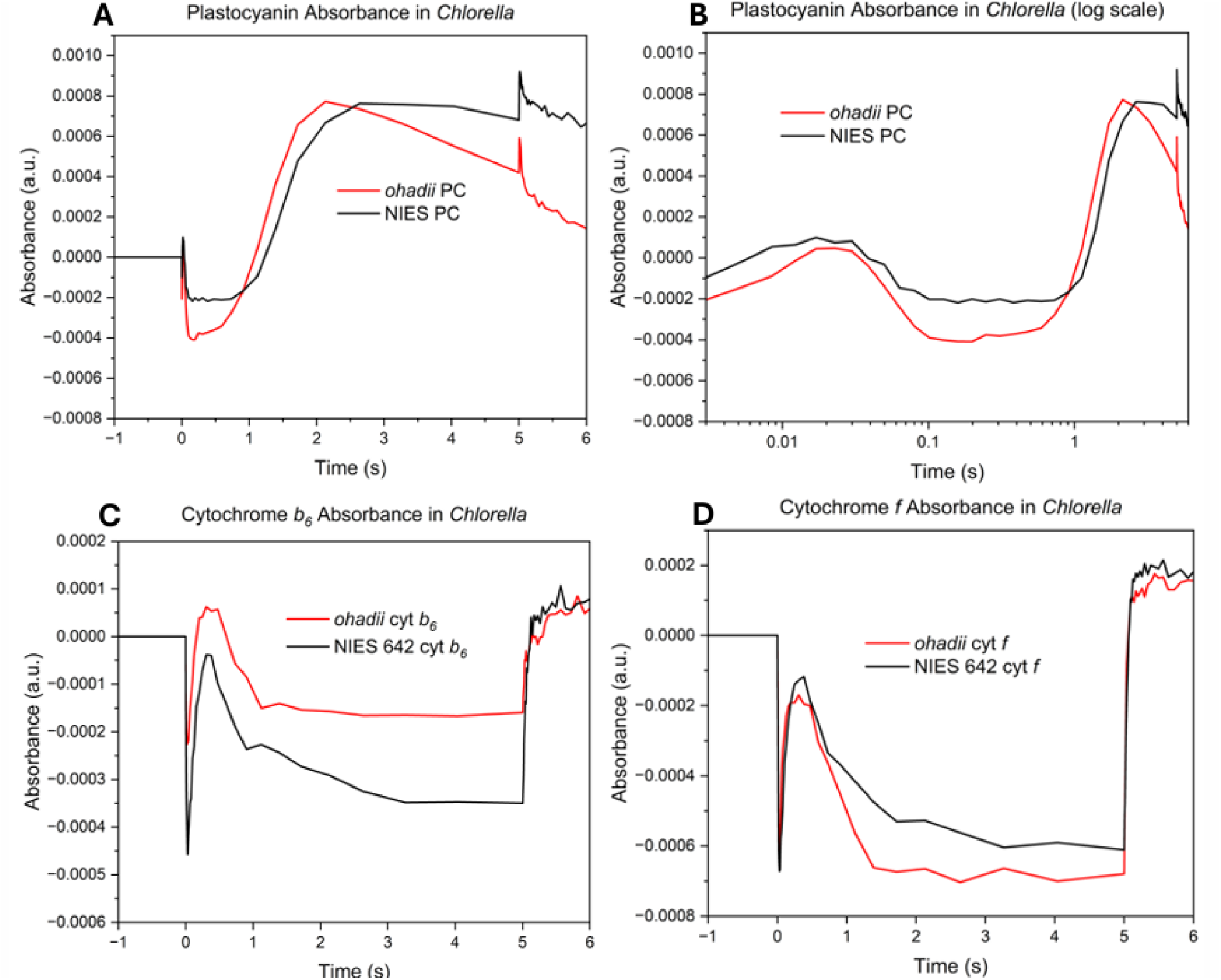
Absorbance change measurements of the cytochrome *b*_*6*_*f* components in *Chlorella*. Absorbance changes over time of the oxidized form of plastocyanin are reported in both **a)** linear and **b)** logarithmic time scales. Absorbance changes due to the reduced forms of **c)** cytochrome *b*_*6*_ and **d)** cytochrome *f* are reported to monitor Q-cycle and linear transfer through cytochrome *b*_*6*_*f*, respectively. The baseline of zero arbitrary units (a.u.) for all panels corresponds to dark absorbance measurement of samples.

The *Chlorella* species display dissimilar amplitudes of both PC and cyt *b*_*6*_ upon both the induction and conclusion of the illumination period (five seconds). In **Fig. 4a**, the overall process of PC redox changes for both organisms appears as an initial reduction of the pool of available PC until approximately one second of illumination. This is followed by rising oxidation to an equivalent absorbance change from the dark baseline (+7.8 × 10^−4^ a.u.), suggesting that expression of PC quantity is independent of PSII-CEF intensity, as both models exhibit the same maximum absorbance of PC at equivalent optical densities. Following this absorbance maximum, *C. ohadii* exhibits a larger slope of PC reduction, likely owing to the accelerated electron transfer to, and continuous utilization of, PQ in the high PSII-CEF system. Comparatively, this amplified reduction of the PC pool in *C. ohadii* occurring while the entirety of the PETC is operating shows that it cannot be the external electron donor side the WOC in the mechanism of PSII-CEF if it is not being continuously oxidized in comparison.

**Fig. 4b** better illustrates electron transfer through PC in time scales applicable to changes caused by charge separation of the two photosystems. At the onset of actinic light, we can now see immediate oxidation of PC by PSI, which occurs faster than the reduction via cyt *b*_*6*_*f* that follows. This occurs as PSI performs charge separation faster than PSII[70], partially due to limitation of photochemistry within the P680 reaction center by LHCII[71]. This slower kinetic of electron transfer through PSII of a few ms[26, 72, 73] coupled with the rate-limitation of PQ diffusion averaging approximately 3-5 ms[2, 62-64] vs PC averaging around 150-550 μs[74, 75], causes the buildup of oxidized PC that begins around one second.

When comparing kinetics for redox states of cyt *b*_*6*_ **(Figure 4c)** and cyt *f* **(Figure 4d)** in tandem, NIES 642 has more equal electron transfer between both cytochrome subunits than *C. ohadii*, with high PSII-CEF resulting in a more reduced cyt *b*_*6*_, and a more oxidized cyt *f*. Inter-species comparison of electron transfer through cyt *f* only slightly differentiates during the equilibrium phase, while cyt *b*_*6*_ amplitudes are substantially different during the entire illumination period. These findings suggest that PSII-CEF does not greatly affect the amplitude of linear flow through cytochrome *b*_*6*_*f*[76], but the process does increase the operation of the Q-cycle attributed with electron flow through cyt *b*_*6*_, where PQH_2_ is reconstituted from electrons donated to cyt *b*_*6*_*f*, driving additional protons from the stroma of the thylakoid to the lumen[17, 68].

### 3.4 77K fluorescence emission

The fluorescence emission spectra of chlorophyll *a* in **Fig. 5** were recorded after flash-freezing whole cells in liquid nitrogen (77K). Reaction center fluorescence emissions at 685 nm (special pair chlorophyll) and 695 nm (trap chlorophyll) were monitored alongside PSI fluorescence emissions that center around 725 nm. By performing Gaussian fits of F_685_, F_695_, and PSI, their relative abundance may be attributed to the area under their fluorescence curve, owing to the dispersion of fluorescent pigments associated with each[51, 77]. Areas beneath the curve, obtained via Gaussian fitting of fluorescent traces, are listed in **Table 3**. Graphical representations of Gaussian fitting for both *Chlorella* species are available in **Supplemental Figure S4**.

**Table 3.** Areas from Gaussian fitting of photosystem components retrieved from the traces in Fig. 5. F_685_ area is a representation of active chlorophyll centers of PSII, combining LHCII, CP47, and CP43 emission. F_695_ area reveals chlorophyll trap usage (CP47).

|  | *C. ohadii* | NIES 642 |
| --- | --- | --- |
| **F<sub>685</sub> Area** | 7430 ± 300 | 9750 ± 500 |
| **F<sub>695</sub> Area** | 1710 ± 310 | 3170 ± 550 |
| **F<sub>685</sub>: F<sub>695</sub>** | 4.34 ± 0.80 | 3.07 ± 0.55 |
| **PSI Complex Area** | 26300 ± 300 | 38100 ± 400 |
| **PSII: PSI** | 0.348 ± 0.017 | 0.340 ± 0.020 |

**Figure 5.**
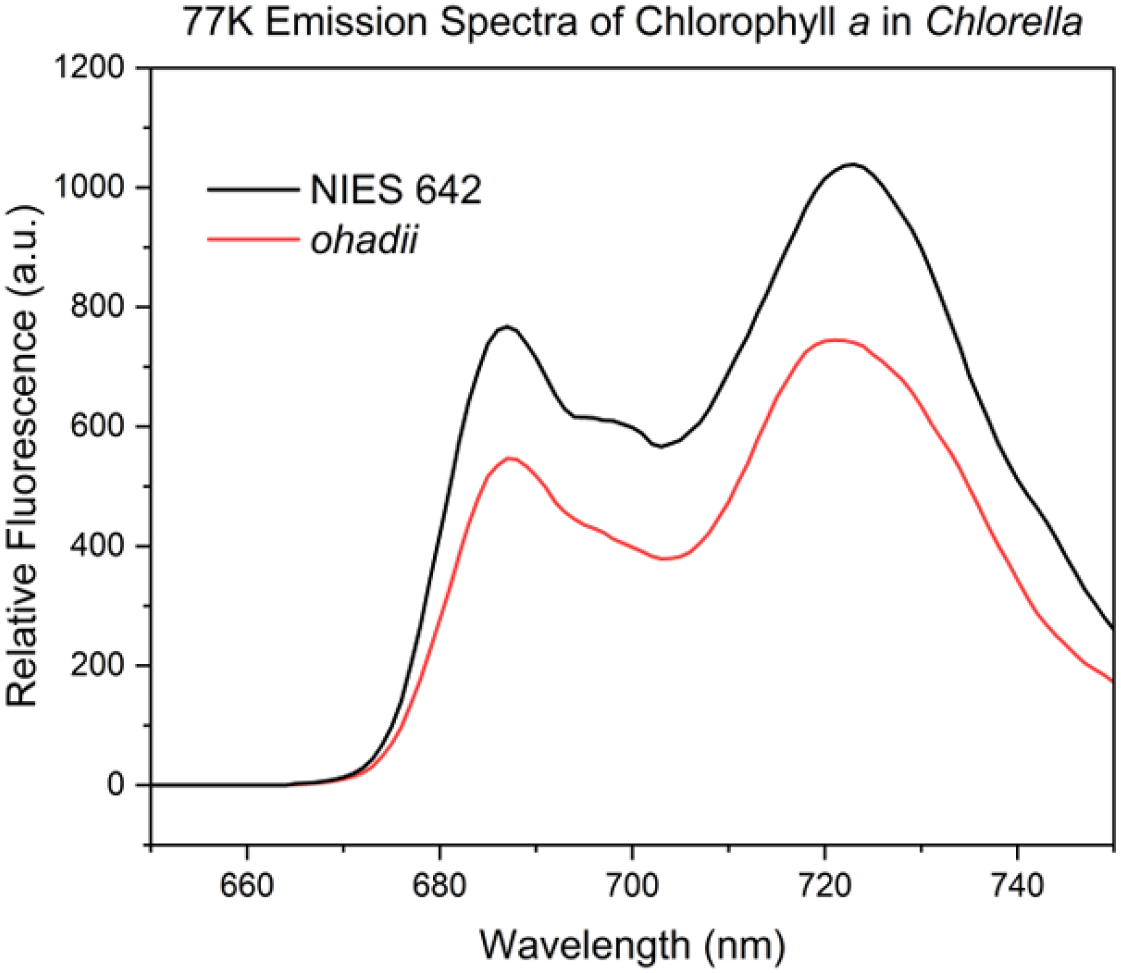
Cryogenic fluorescence emission spectra from excitation of chlorophyll *a* at 435 nm in flash-frozen *Chlorella* cells. Spectra are normalized to OD_730_.

The higher ratio of reaction center to trap chlorophyll in *C. ohadii* (4.34 versus 3.07 in NIES 642) demonstrates the exceptional photoprotective abilities of the organism. This minimal fluorescence broadening is present, even after several generations of growth at an irradiance level that would induce irreparable photoinhibition in most phototrophic organisms within a few hours[78, 79]. The ratio of pigmentation across photosystems is conserved between species, with a difference of just over 2%. The lack of a significant increase in PSII:PSI ratio in *C. ohadii* is unusual, as higher light conditions frequently result in a down regulation of PSI concentrations in phototrophic organisms[80, 81]. Further inter-species comparison of PSI was therefore evaluated via observations of kinetics of the P700 reaction center, reported in **Fig. 6**.

**Figure 6.**
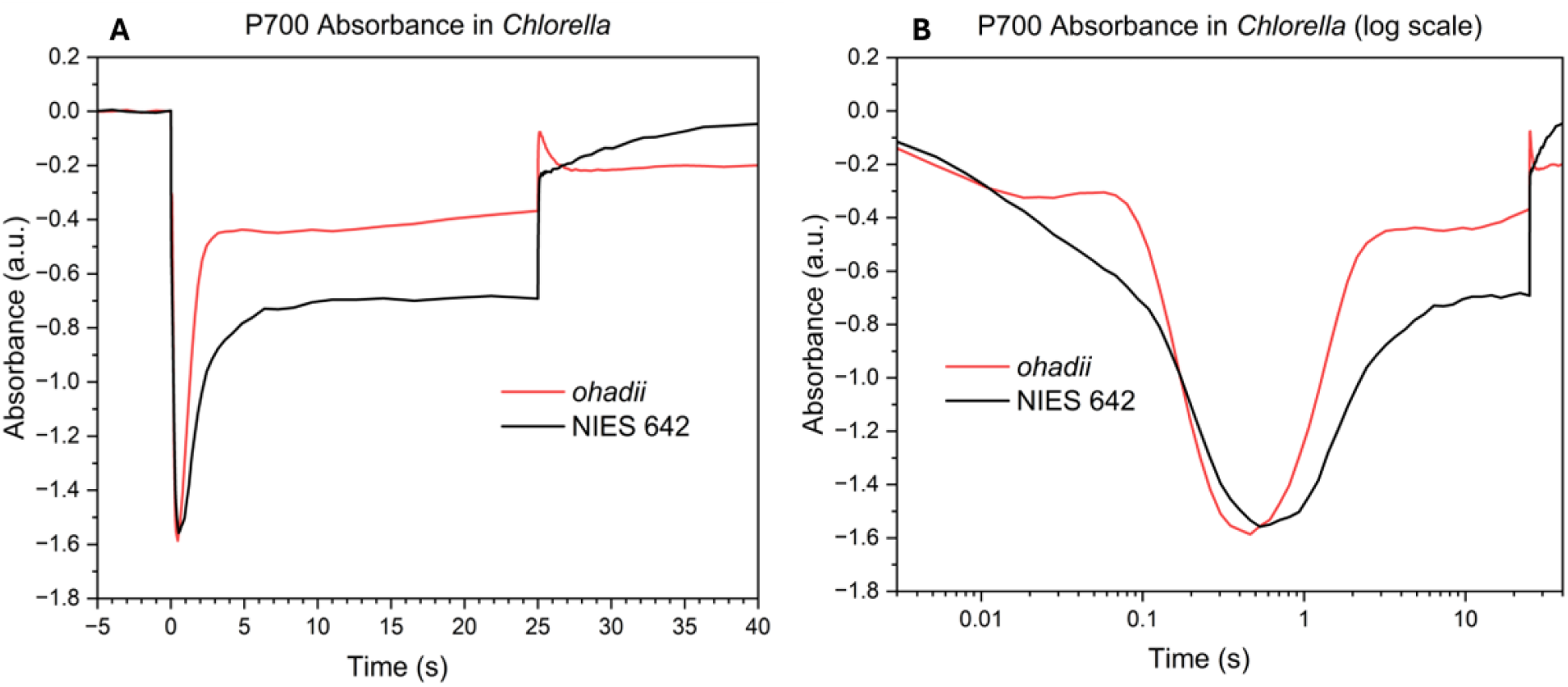
Redox kinetics of the photosystem I reaction center, P700, measured by absorbance changes at 810 nm in both **a)** linear and **b)** logarithmic timescales. Photon capture via transfer from antenna chlorophyll surrounding PSI causes a decrease in absorbance as P700 performs charge separation to the A_0_ chlorophyll.

### 3.5 P700 absorbance

Kinetics and amplitude of the PSI reaction center were monitored by absorbance changes under actinic illumination. Linear timescale traces of P700 absorbance of both *Chlorella* species are illustrated in **Figure 6a** as an overall view of PSI amplitude, with logarithmic timescales for observation of ms timescale kinetics in **Figure 6b**.

As charge separation events begin at the onset of light, there is a notable pause in the overall oxidation of PSI in *C. ohadii*, while NIES 642 has a more linear increase to its maximum operation. Both species reach a fairly equal maximum oxidation of PSI; however, *C. ohadii* reaches this maximum hundreds of ms faster despite the initial lag in operation. Around 400-500 ms the rate of chemical energy leaving the PETC is outpaced by the reducing energy created by the photosystems, causing a buildup of reduced PSI. In both species, the redox poise of P700 quickly equilibrates, with *C. ohadii* reaching a steady-state far closer to the dark baseline. Based on the equal portioning of photosystem stoichiometry presented in **Table 3** and the equivalent maximum absorbance change in **Fig. 6**, both species have approximately equal quantity and activity of PSI. Despite this, *C. ohadii* shows a significantly higher quantity of reduced PSI at steady-state equilibrium.

When the light is shut off, NIES 642 consistently re-reduces the PSI centers via upstream electron transfer, but *C. ohadii* exhibits a distinctly separate morphology. Immediately, there is a stark return towards the baseline level of reduced PSI in the red trace. What appears to be a sizable delivery of reductant to P700 in the absence of light may be the PQ pool of *C. ohadii*, burdened with carrying, and likely constantly recycling, the excess reducing power leaving PSII. The red trace also does not return to its original dark baseline, suggesting a redox change in the overall PSI population of *C. ohadii* during illumination. An increase in PSI-CEF by the existence of cyt *b*_*6*_*f*-PSI supercomplexes may explain the initial wave of quick reduction to PSI (<100 ms) and the lower redox change from dark to light levels in *C. ohadii*[82, 83]. PSI-CEF may be used in tandem with PSII-CEF as high-light conditions are introduced and downstream reactions need to be expedited.

### 3.6 Electrochromic shift

Electrochromic band-shift of carotenoids within PSI was quantified under actinic light activation as a means of measuring the amplitude of proton motive force (pmf) changes in *Chlorella* cells (**Fig. 7)**. The pmf is established by two main contributing factors, the trans-thylakoid membrane potential (ΔΨ) and the proton concentration gradient (ΔpH)[84, 85]. In various photosynthetic organisms, the pmf has been shown to modulate the rate of the PETC from PSII to PSI, and its modulation is imperative for the survival of phototrophs experiencing increasing light conditions[63, 86, 87]. Analyzing the log-scale kinetics of ECS in **Figure 7b** reveals the amplitudes of processes within the PETC that regulate proton transfer across the thylakoid.

**Figure 7.**
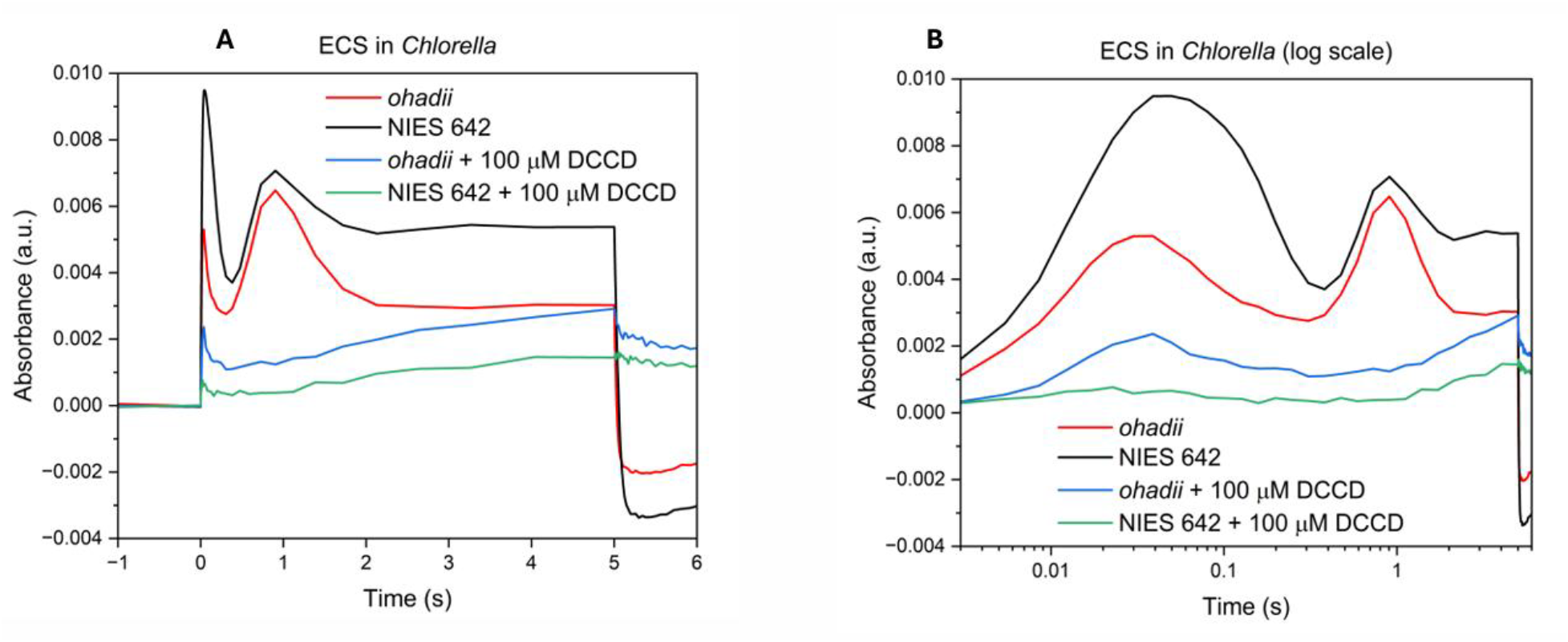
Electrochromic band-shift (ECS) spectra of both *Chlorella* strains. Linear timescale ECS traces are shown in panel **a)**, while log-scale kinetics are presented in **b)**. Positive (upward) changes from baseline are attributed to the ΔΨ component, membrane potential, and negative (downward) changes are caused by the proton concentration gradient, ΔpH. 100 μM DCCD inhibited samples were prepared in darkness to avoid PETC operation prior to testing.

N,N’-dicyclohexylcarbodiimide (DCCD) is a common PETC inhibitor that can be useful in ECS, as it diminishes the ΔpH component of the pmf[88, 89]. DCCD works by covalently binding to the F0 subunit of ATP-synthase, preventing the rotating mechanism of the enzyme that causes proton efflux from the lumen of the thylakoid[90, 91]. 100 μM concentrations of DCCD were used to inhibit the contributions of ATP-synthase in additional traces (blue and green) of **Fig. 7**.

The rise in ECS in the first several ms stems from the charge separating events and Q-cycle of the PETC causing a rise in ΔΨ, and proton deposition into the luminal space. This concentration gradient is leveraged by ATP-synthase operation, causing a fall in ECS due to the change in pH. The second rise that follows is attributed to a slower relaxation of the ΔpH component of the pmf[92]. After steady-state is reached as the carbon fixation reactions equilibrate the rates of the PETC and ATP synthase operation, the amplitude of the trace’s fall below the baseline grants insight to relative ATP synthase distribution by isolation of the contribution of ΔpH.

NIES 642 quickly establishes a strong potential across the membrane and maintains a stronger ΔΨ than *C. ohadii* for the entirety of the PETC operation. With initial proton deposition into the lumen being facilitated by the WOC removing hydrogen from water, the higher oscillation quality in low PSII-CEF would in turn create a greater concentration gradient. *C. ohadii* then exhibits a slightly faster rebound back toward the baseline potential, exhibiting an expedited ATP-synthase operation, parallel to what was seen for *C. ohadii* PSI. Once the machinery of carbon fixation is activated, the trans-thylakoid potential is raised again in both species as ATP is consumed (∼300 ms). Once this carbon fixation reaches maximum turnover velocity, ATP synthase kinetics equilibrate with carbon fixation kinetics around two seconds in both models. For the duration of the illumination, *C. ohadii* has a proton gradient closer to its gradient in the dark.

In the traces with the presence of DCCD, minimal changes of membrane potential and pH are present in both cultures over the entire period. The lack of electrostatic membrane changes surprisingly assert that photosystem operation has been halted (barring a small resistance in the initial rise of the *C. ohadii* trace). Computationally, the linear rise in ECS over the illumination stems from the gradual disappearance of the corrective measurement of light scattering through the thylakoid (546 nm). Previously, DCCD has been shown to bind to the luminal PSII subunit PsbS[93, 94], and this association would eliminate a second major component of the heat dissipating mechanism of most green algae, potentially causing diffusion of water into the thylakoid due to the high proton gradient within the lumen.

## 4. Discussion

Broad overview of the two PSII-CEF models shows that these *Chlorella* species have numerous differences in the operation of their PETC that have arisen from exponentially different needs of photoprotection. The first major consequence of this mechanism is that it diminishes the efficiency of the PSII donor side in exchange for retaining the reducing potential captured from light within the PETC (minimizing reactive oxygen species). This is to say that, while NIES 642 receives 100 times fewer incident photons naturally, these photons are being utilized more effectively for linear electron flow and water splitting, causing a higher quantum yield in the low PSII-CEF model when monitored by rate oximetry. Analysis of donor side efficiency has proven advantageous for comparatively monitoring the presence of PSII-CEF, detailing an absence of the backwards transitions in NIES 642. Despite the ability of FRR to verify and quantify electron donation to the WOC, it does not differentiate between the possible intermediates performing this PSII-CEF. To identify the electron carrier(s) in the mechanism of PSII-CEF, attention shifts towards the operation and regulation of the acceptor side, where PQ likely plays some form of regulatory role.

The speed with which PSI begins operation, the level it is re-reduced by PC, and the consistent reduction of PC in *C. ohadii* during continuous illumination collectively assert that PC cannot be the redox carrier back to the WOC in the mechanism of PSII-CEF. The rate-limiting step at the Q-cycle’s cyt *b*_*6*_*f* end is probably driving the PQ pool to a more reduced equilibrium. We can also see from 77K fluorescence emission that *C. ohadii* expresses a higher ratio of reaction center to chlorophyll trap pigmentation compared to NIES 642 (4.34 to 3.07). This minimal fluorescence broadening towards the core antennae of PSII (F_695_) speaks toward the efficiency of photoprotection in *C. ohadii* cells. These observations collectively strongly suggest two mechanistic insights. First, the electrons being placed onto quinone to form semiquinone in Q_B_ are likely equilibrating with an internal acceptor, putatively (semi)quinone in Q_C_, as a replacement for the chlorophyll trap in the cause of over-excitation. Second, if there is an external carrier returning electrons to the WOC, it must be PQH_2_ and not PC.

One theorized secondary cofactor of electron acceptance from Q_B_ in PSII-CEF is the heterodimeric cyt *b*_*559*,_ made of α (PsbE) and β (PsbF) subunits[60, 95]. At an approximate distance of 25 Å from the Q_B_ site in PSII and relatively stromally positioned, cyt *b*_*559*_ is likely not the primary electron acceptor in the PSII-CEF pathway from Q_B_ directly, especially in organisms with high levels of PSII-CEF where ms timescales of electron transfer would be necessary for adequate photoprotection[32]. However, a plausible schematic for electron transfer to the closer, putative Q_C_ site was recently reported, which hypothesized backward transitions of the WOC being contingent on reduction of the acceptor pool and (semi)quinone Q_C_[23]. The findings listed in **Table 3** corroborate this hypothesis, showing that in the high PSII-CEF species the PQ pool is being almost continuously turned over and quickly reduced by Q_A_^-^. Meanwhile, there is a proportionally low (38.7%) amount of PSII with associated semiquinone in the Q_B_ site, despite the semiquinone form commonly exhibiting a more stable association with PSII, as observed in NIES 642. Considering most Q_B_ sites are occupied in the high PSII-CEF system, yet most of the associated PQ is not the more stable semiquinone form[96-98], it is reasonable that there may be a substantial amount of PQ bound within the Q_C_ site, with the semiquinone form equilibrating between Q_B_ and Q_C_. Coupling this pathway to an external mechanism utilizing PQH_2_ for electron transfer to the WOC also stays in line with the well-supported theme that various alternate electron pathways of the PETC help to regulate the energetic needs of the organism by prioritizing the production of ATP[1, 7-9, 82, 99] via increased proton transport into the lumen.

The ratio of chemical energy made from photosynthesis would be perfectly balanced for carbon fixation itself, but ATP is utilized for several other necessary processes in the phototroph other than powering RuBisCO[100]. As evidenced in **Figure 7**, the proton gradient of *C. ohadii* remains closer throughout the entirety of ECS measurements to its proton gradient in the dark, maintaining about half the amplitude of the NIES 642 trace, both initially and at steady-state. This smaller gradient and shallower drop in ECS at the termination of the light speaks to the strength of ATP synthase activity in the high PSII-CEF model, especially if the enhanced removal of protons from the stroma by the amplified Q-cycle is taken into account. The role of Q_C_ in this mechanism would therefore be a rapid, powerful safety valve capable of ensuring electrons are removed from semiquinone in the Q_B_ site on a timescale which will both prevent damaging recombination to triplet chlorophyll[101] and allow electrons to leave the photosystem when conditions are favorable. This timescale is probably similar to the frequency of excitons received, i.e. hundreds of microseconds per reaction center excitation[23, 32, 35, 36].

To achieve an oxygen evolution rate of 1,254 μmol O_2_/mg chl *a*/h, *C. ohadii* must also perform a considerable amount of linear electron flow, regardless of the ratio of LEF:PSII-CEF. For context of typical phototrophic organisms, *Synechocystis* sp. PCC 6803, a model cyanobacterium, produces between 270-329 μmol O_2_/mg chl *a*/h when illuminated with 500 µEin/m^2^/s[102]. Due to the high solar radiation environment of the Negev, with minimal shading during its natural light cycle, the optimization of photosystems seems inherently necessary for *C. ohadii* to prioritize sequestering the products of the PETC promptly, due to the rate limitations of photosynthesis being dependent on RuBisCO activity (carbon fixation)[103]. Literature regarding the isolation of *C. ohadii* has observed that this species possesses a large pyrenoid, enabling a 10-fold increase in association of inorganic carbon via the induction of a carbon concentration mechanism in mixotrophic conditions[40].

ATP synthase and PSI have seemingly optimized in conjunction to utilize the balance of electron transfer from PSII in *C. ohadii*. Increasing the stoichiometric ratio of the two photosystems towards PSI has been shown to optimize ATP production via PSI-CEF[99], and it stands to reason that these two photoprotective mechanisms would be employed in tandem. Redox poise changes at PSI in the dark on the timescale of seconds, along with increases of reduction to PC and PSI compared to NIES 642 point towards modulation of the photosystem into a supercomplex, increasing the ratio of ATP:NADPH. Unfortunately, a comprehensive crystal structure of either photosynthetic apparatus at the natural light conditions of *C. ohadii* is not currently available for study. The authors hope this issue may be addressed in further work following the results herein.

## 5 Conclusions

The model of high PSII-CEF, *C. ohadii*, shows an increase in each common inefficiency parameter of the WOC when compared to low PSII-CEF NIES 642. This difference in WOC efficiency results in an order of magnitude difference in the quantum efficiency of these species. At extreme light conditions, *C. ohadii* performs an almost continuous turnover of PQ that employs several downstream consequences in the photosynthetic apparatus. Over-reduction of the PQ pool diminishes ΔΨ due to the overabundance of negative charge on the luminal side of the PETC and increases the ΔpH by up-regulating the Q-cycle. Seemingly via adaptations to an increased proton influx to the lumen of the thylakoid, *C. ohadii* establishes a proton gradient much closer to its dark baseline by increasing the operation of ATP synthase. The operation of this protein, along with PSI operation, is expedited in response, likely due to a quick enactment of PSI-CEF. This increase in CEF around both photosystems protects the PETC by regulating its redox poise, while increasing growth of the organism by the optimization of ATP production.

## Acknowledgments

The group would like to thank the following individuals for their contributions towards our comparative experiments: Dr. David Vinyard for the development of the VZAD software program, Dr. Debashish Bhattacharya for valuable insights, and Dr. Michael Vaughn for the continued development of the JTS-150 spectrofluorometer.

## Data Availability

Data will be made available on request. For additional information regarding the contents of this article, please contact the corresponding author.

## CRediT Author Contribution Statement

**Grant Steiner:** Validation, Formal Analysis, Investigation, Writing-Original Draft, Visualization. **Devinjeet Saini:** Software, Investigation **Arivu Kapoor:** Investigation. **Colin Gates:** Conceptualization, Methodology, Software, Writing-Review & Editing, Visualization, Supervision.

## References

1. Farooq, M., et al., Drought Stress in Plants: An Overview, in Plant Responses to Drought Stress: From Morphological to Molecular Features, R. Aroca, Editor. 2012, Springer Berlin Heidelberg: Berlin, Heidelberg. p. 1–33.

2. Rochaix, J.-D., Regulation of photosynthetic electron transport. Biochimica et Biophysica Acta (BBA) - Bioenergetics, 2011. 1807(3): p. 375–383.

3. Chadee, A., et al., The Complementary Roles of Chloroplast Cyclic Electron Transport and Mitochondrial Alternative Oxidase to Ensure Photosynthetic Performance. Frontiers in Plant Science, 2021. 12.

4. Ruban, A.V. and S. Wilson, The Mechanism of Non-Photochemical Quenching in Plants: Localization and Driving Forces. Plant and Cell Physiology, 2020. 62(7): p. 1063–1072.

5. Allen, J.F., Photosynthesis of ATP—Electrons, Proton Pumps, Rotors, and Poise. Cell, 2002. 110(3): p. 273–276.

6. Zhang, S., et al., Structural insights into photosynthetic cyclic electron transport. Molecular Plant, 2023. 16(1): p. 187–205.

7. Munekage, Y., et al., Cyclic electron flow around photosystem I is essential for photosynthesis. Nature, 2004. 429(6991): p. 579–582.

8. Joliot, P., et al., High efficient cyclic electron flow and functional supercomplexes in Chlamydomonas cells. Biochimica et Biophysica Acta (BBA) - Bioenergetics, 2022. 1863(8): p. 148909.

9. Shikanai, T. and H. Yamamoto, Contribution of Cyclic and Pseudo-cyclic Electron Transport to the Formation of Proton Motive Force in Chloroplasts. Molecular Plant, 2017. 10(1): p. 20–29.

10. Kuntz, M., Plastid terminal oxidase and its biological significance. Planta, 2004. 218(6): p. 896–899.

11. McDonald, A.E., et al., Flexibility in photosynthetic electron transport: The physiological role of plastoquinol terminal oxidase (PTOX). Biochimica et Biophysica Acta (BBA) - Bioenergetics, 2011. 1807(8): p. 954–967.

12. Falkowski, P.G., et al., Evidence for Cyclic Electron Flow around Photosystem II in Chlorella pyrenoidosa. Plant Physiol., 1986. 81: p. 310–312.

13. Mullineaux, P.M., et al., ROS-dependent signalling pathways in plants and algae exposed to high light: Comparisons with other eukaryotes. Free Radical Biology and Medicine, 2018. 122: p. 52–64.

14. Murata, N., et al., Photoinhibition of photosystem II under environmental stress. Biochimica et Biophysica Acta, 2007. 1767: p. 414–421.

15. Ananyev, G., C. Gates, and G.C. Dismukes, The Oxygen quantum yield in diverse algae and cyanobacteria is controlled by partitioning of flux between linear and cyclic electron flow within photosystem II. Biochimica et Biophysica Acta, 2016. 1857: p. 1380–1391.

16. Van Eerden, F.J., et al., Exchange pathways of plastoquinone and plastoquinol in the photosystem II complex. Nature Communications, 2017. 8(1): p. 15214.

17. Cramer, W.A., S.S. Hasan, and E. Yamashita, The Q cycle of cytochrome bc complexes: A structure perspective. Biochimica et Biophysica Acta (BBA) - Bioenergetics, 2011. 1807(7): p. 788–802.

18. Kamada, S., Y. Nakajima, and J.-R. Shen, Structural insights into the action mechanisms of artificial electron acceptors in photosystem II. Journal of Biological Chemistry, 2023. 299(7).

19. Umena, Y., et al., Crystal structure of oxygen-evolving photosystem II at a resolution of 1.9 Å. Nature, 2011: p. 1–7.

20. Hasegawa, K. and T. Noguchi, Molecular interactions of the quinone electron acceptors QA, QB, and QC in photosystem II as studied by the fragment molecular orbital method. Photosynthesis Research, 2014. 120(1): p. 113–123.

21. Kaminskaya, O., V.A. Shuvalov, and G. Renger, Evidence for a Novel Quinone-Binding Site in the Photosystem II (PS II) Complex That Regulates the Redox Potential of Cytochrome b559. Biochemistry, 2007. 46(4): p. 1091–1105.

22. Kedem, I., et al., Juggling Lightning: How Chlorella ohadii handles extreme energy inputs without damage. Photosynthesis Research, 2021. 147(3): p. 329–344.

23. Gates, C., et al., Regulation of light energy conversion between linear and cyclic electron flow within photosystem II controlled by the plastoquinone/quinol redox poise. Photosynth Res, 2023. 156(1): p. 113–128.

24. Linke, K. and F.M. Ho, Water in Photosystem II: Structural, functional and mechanistic considerations. Biochimica et Biophysica Acta (BBA) - Bioenergetics, 2014. 1837(1): p. 14–32.

25. Doyle, M.D., et al., Water Networks in Photosystem II Using Crystalline Molecular Dynamics Simulations and Room-Temperature XFEL Serial Crystallography. Journal of the American Chemical Society, 2023. 145(27): p. 14621–14635.

26. Vinyard, D.J., G. Ananyev, and G.C. Dismukes, Photosystem II: The Reaction Center of Oxygenic Photosynthesis. Annual Review of Biochemistry, 2013. 82(1): p. 577–606.

27. Baker, N.R., J. Harbinson, and D.M. Kramer, Determining the limitations and regulation of photosynthetic energy transduction in leaves. Plant, Cell & Environment, 2007. 30(9): p. 1107–1125.

28. Pantazis, D.A., Missing Pieces in the Puzzle of Biological Water Oxidation. ACS Catalysis, 2018. 8(10): p. 9477–9507.

29. Kok, B., B. Forbush, and M. McGloin, Cooperation of charges in photosynthetic O2 evolution-I. A linear four step mechanism. Photochemistry and Photobiology, 1970. 11(6): p. 457–475.

30. Ishikita, H. and K. Saito, Photosystem II: Probing Protons and Breaking Barriers. Biochemistry, 2025.

31. Messinger, J., W.P. Schroeder, and G. Renger, Structure-function relations in photosystem II. Effects of temperature and chaotropic agents on the period four oscillation of flash-induced oxygen evolution. Biochemistry, 1993. 32(30): p. 7658–7668.

32. Ananyev, G., et al., Photosystem II-cyclic electron flow powers exceptional photoprotection and record growth in the microalga Chlorella ohadii. Biochimica et Biophysica Acta (BBA) - Bioenergetics, 2017. 1858(11): p. 873–883.

33. Kern, J., et al., Structures of the intermediates of Kok’s photosynthetic water oxidation clock. Nature, 2018. 563(7731): p. 421–425.

34. Mino, H. and A. Kawamori, EPR studies of the water oxidizing complex in the S1 and the higher S states: the manganese cluster and YZ radical. Biochimica et Biophysica Acta (BBA) - Bioenergetics, 2001. 1503(1): p. 112–122.

35. Ananyev, G. and G. Dismukes, How fast can photosystem II split water? Kinetic performance at high and low frequencies. Photosynthesis Research, 2005.

36. Gates, C., G. Ananyev, and G.C. Dismukes, Realtime kinetics of the light driven steps of photosynthetic water oxidation in living organisms by “stroboscopic” fluorometry. Biochimica et Biophysica Acta (BBA) - Bioenergetics, 2020. 1861(8): p. 148212.

37. Vinyard, D.J., et al., Thermodynamically accurate modeling of the catalytic cycle of photosynthetic oxygen evolution: A mathematical solution to asymmetric Markov chains. Biochim Biophys Acta, 2013. 1827(7): p. 861–8.

38. Lewis, L.A. and R.M. McCourt, Green algae and the origin of land plants. American Journal of Botany, 2004. 91(10): p. 1535–1556.

39. Metsoviti, M.N., et al. Effect of Light Intensity and Quality on Growth Rate and Composition of Chlorella vulgaris. Plants, 2020. 9, DOI: 10.3390/plants9010031.

40. Treves, H., et al., The mechanisms whereby the green alga Chlorella ohadii, isolated from desert soil crust, exhibits unparalleled photodamage resistance. New Phytologist, 2016.

41. Treves, H., et al., A newly isolated Chlorella sp. from desert sand crusts exhibits a unique resistance to excess light intensity. FEMS Microbiology Ecology, 2013. 86(3): p. 373–380.

42. Fadeeva, M., et al. Structure of Chlorella ohadii Photosystem II Reveals Protective Mechanisms against Environmental Stress. Cells, 2023. 12, DOI: 10.3390/cells12151971.

43. Takamura, N., F. Kasai, and M. Watanabe, Differences in the tolerant level of benthic algae to heavy metal-The effects of Cu, Cd, and Zn on the photosynthesis. Res. Rep. Natl. Inst. Environ. Stud, 1988(114): p. 223–232.

44. Dall’Osto, L., et al., Combined resistance to oxidative stress and reduced antenna size enhance light-to-biomass conversion efficiency in Chlorella vulgaris cultures. Biotechnology for Biofuels, 2019. 12(1): p. 221.

45. Al-Mayah, A. and M. Naeemah, Cultivation of Chlorella Vulgaris in BG-11 Media Using Taguchi Method. Journal of Advanced Research in Dynamical and Control Systems, 2018. 10: p. 19–30.

46. Stanier, R.Y., et al., Purification and properties of unicellular blue-green algae (order Chroococcales). Bacteriological reviews, 1971. 35(2): p. 171–205.

47. Gorbunov, M.Y., Z.S. Kolber, and P.G. Falkowski, Measuring photosynthetic parameters in individual algal cells by Fast Repetition Rate fluorometry. Photosynthesis Research, 1999. 62(2): p. 141–153.

48. Suggett, D.J., O. Prášil, and M.A. Borowitzka, Chlorophyll a fluorescence in aquatic sciences: methods and applications. Vol. 4. 2010: Springer.

49. Delieu, T. and D.A. Walker, AN IMPROVED CATHODE FOR THE MEASUREMENT OF PHOTOSYNTHETIC OXYGEN EVOLUTION BY ISOLATED CHLOROPLASTS. New Phytologist, 1972. 71(2): p. 201–225.

50. Porra, R. and W. Thompson, Determination of accurate extinction coefficients and simultaneous equations for assaying chlorophylls a and b extracted with four different solvents: verification of the …. Biochim Biophys Acta, 1989.

51. Lamb, J.J., G. Røkke, and M.F. Hohmann-Marriott, Chlorophyll fluorescence emission spectroscopy of oxygenic organisms at 77 K. Photosynthetica, 2018. 56(1): p. 105–124.

52. Harbinson, J. and C.L. Hedley, Changes in P-700 Oxidation during the Early Stages of the Induction of Photosynthesis. Plant Physiology, 1993. 103(2): p. 649–660.

53. Mathiot, C. and J. Alric, Standard units for ElectroChromic Shift measurements in plant biology. Journal of Experimental Botany, 2021. 72(18): p. 6467–6473.

54. Kramer, D.M. and C.A. Sacksteder, A diffused-optics flash kinetic spectrophotometer (DOFS) for measurements of absorbance changes in intact plants in the steady-state. Photosynthesis Research, 1998. 56(1): p. 103–112.

55. Sane, P.V., et al., A Study of State Changes in Chlorella: The Effect of Uncoupler and Energy Transfer Inhibitors. Zeitschrift für Naturforschung C, 1982. 37(5-6): p. 458–463.

56. Shinkarev, V.P., Flash-Induced Oxygen Evolution in Photosynthesis: Simple Solution for the Extended <em>S</em>-State Model that includes Misses, Double-Hits, Inactivation, and Backward-Transitions. Biophysical Journal, 2005. 88(1): p. 412–421.

57. Wang, S., D. Zhang, and X. Pan, Effects of arsenic on growth and photosystem II (PSII) activity of Microcystis aeruginosa. Ecotoxicology and Environmental Safety, 2012. 84: p. 104–111.

58. Küpper, H., et al., Analysis of OJIP Chlorophyll Fluorescence Kinetics and QA Reoxidation Kinetics by Direct Fast Imaging. Plant Physiology, 2018. 179(2): p. 369–381.

59. Zournas, A., K. Mani, and G.C. Dismukes, Cyclic electron flow around photosystem II in silico: How it works and functions in vivo. Photosynthesis Research, 2023. 156(1): p. 129–145.

60. Shinopoulos, K.E. and G.W. Brudvig, Cytochrome b 559 and cyclic electron transfer within photosystem II. Biochimica et Biophysica Acta (BBA)-Bioenergetics, 2012. 1817(1): p. 66–75.

61. Onno Feikema, W., et al., Cyclic electron transfer in photosystem II in the marine diatom Phaeodactylum tricornutum. Biochimica et Biophysica Acta (BBA) - Bioenergetics, 2006. 1757(7): p. 829–834.

62. Robinson, H.H. and A.R. Crofts, Kinetics of the oxidation—reduction reactions of the photosystem II quinone acceptor complex, and the pathway for deactivation. FEBS Letters, 1983. 153(1): p. 221–226.

63. Tikhonov, A.N., The cytochrome b6f complex at the crossroad of photosynthetic electron transport pathways. Plant Physiology and Biochemistry, 2014. 81: p. 163–183.

64. Kirchhoff, H., U. Mukherjee, and H.J. Galla, Molecular Architecture of the Thylakoid Membrane: Lipid Diffusion Space for Plastoquinone. Biochemistry, 2002. 41(15): p. 4872–4882.

65. Gates, C., et al., Exceptional Quantum Efficiency Powers Biomass Production in Halotolerant Algae Picochlorum sp. Photosynthesis research, 2024: p. 1–19.

66. Beauchemin, R., et al., Spermine and spermidine inhibition of photosystem II: Disassembly of the oxygen evolving complex and consequent perturbation in electron donation from TyrZ to P680+ and the quinone acceptors QA− to QB. Biochimica et Biophysica Acta (BBA) - Bioenergetics, 2007. 1767(7): p. 905–912.

67. Torzillo, G., et al., Interplay between photochemical activities and pigment composition in an outdoor culture of Haematococcus pluvialis during the shift from the green to red stage. Journal of Applied Phycology, 2003. 15(2): p. 127–136.

68. Baniulis, D., et al., Structure–Function of the Cytochrome b6f Complex. Photochemistry and Photobiology, 2008. 84(6): p. 1349–1358.

69. Datta, S.N., J. Sudhamsu, and A. Pandey, Theoretical Determination of the Standard Reduction Potential of Plastocyanin in Vitro. The Journal of Physical Chemistry B, 2004. 108(23): p. 8007–8016.

70. Caffarri, S., et al., A comparison between plant photosystem I and photosystem II architecture and functioning. Current Protein and Peptide Science, 2014. 15(4): p. 296–331.

71. Jennings, R.C., et al., Selective quenching of the fluorescence of core chlorophyll–protein complexes by photochemistry indicates that Photosystem II is partly diffusion limited. Photosynthesis Research, 2000. 66(3): p. 225–233.

72. Rutherford, A.W., Photosystem II, the water-splitting enzyme. Trends in Biochemical Sciences, 1989. 14(6): p. 227–232.

73. Ausländer, W. and W. Junge, The electric generator in the photosynthesis of green plants. II. Kinetic correlation between protolytic reactions and redox reactions. Biochimica et Biophysica Acta (BBA) - Bioenergetics, 1974. 357(2): p. 285–298.

74. Hippler, M., et al., Fast Electron Transfer from Cytochrome c6 and Plastocyanin to Photosystem I of Chlamydomonas reinhardtii Requires PsaF. Biochemistry, 1997. 36(21): p. 6343–6349.

75. Höhner, R., et al., Plastocyanin is the long-range electron carrier between photosystem II and photosystem I in plants. Proceedings of the National Academy of Sciences, 2020. 117(26): p. 15354–15362.

76. Malone, L.A., et al., Cytochrome b6f – Orchestrator of photosynthetic electron transfer. Biochimica et Biophysica Acta (BBA) - Bioenergetics, 2021. 1862(5): p. 148380.

77. Busheva, M., A. Andreeva, and E. Apostolova, Effect of modification of light-harvesting complex II on fluorescence properties of thylakoid membranes of Arabidopsis thaliana. Journal of Photochemistry and Photobiology B: Biology, 2000. 56(1): p. 78–84.

78. Tyystjärvi, E. and E.M. Aro, The rate constant of photoinhibition, measured in lincomycin-treated leaves, is directly proportional to light intensity. Proceedings of the National Academy of Sciences, 1996. 93(5): p. 2213–2218.

79. Allakhverdiev, S.I. and N. Murata, Environmental stress inhibits the synthesis de novo of proteins involved in the photodamage–repair cycle of Photosystem II in Synechocystis sp. PCC 6803. Biochimica et Biophysica Acta (BBA) - Bioenergetics, 2004. 1657(1): p. 23–32.

80. Hihara, Y. and M. Ikeuchi, Toward the Elucidation of Physiological Significance of pmgA- mediated High-Light Acclimation to Adjust Photosystem Stoichiometry: Effects of the Prolonged High-Light Treatment on pmgA Mutants, in Photosynthesis: Mechanisms and Effects: Volume I– V: Proceedings of the XIth International Congress on Photosynthesis, Budapest, Hungary, August 17–22, 1998, G. Garab, Editor. 1998, Springer Netherlands: Dordrecht. p. 2929–2932.

81. Sonoike, K., Y. Hihara, and M. Ikeuchi, Physiological Significance of the Regulation of Photosystem Stoichiometry upon High Light Acclimation of Synechocystis sp. PCC 6803. Plant and Cell Physiology, 2001. 42(4): p. 379–384.

82. Iwai, M., et al., Isolation of the elusive supercomplex that drives cyclic electron flow in photosynthesis. Nature, 2010. 464(7292): p. 1210–1213.

83. Zivcak, M., et al., Photosynthetic proton and electron transport in wheat leaves under prolonged moderate drought stress. Journal of Photochemistry and Photobiology B: Biology, 2014. 137: p. 107–115.

84. Armbruster, U., et al., The regulation of the chloroplast proton motive force plays a key role for photosynthesis in fluctuating light. Current Opinion in Plant Biology, 2017. 37: p. 56–62.

85. Kramer, D.M., J.A. Cruz, and A. Kanazawa, Balancing the central roles of the thylakoid proton gradient. Trends in Plant Science, 2003. 8(1): p. 27–32.

86. Armbruster, U., et al., Ion antiport accelerates photosynthetic acclimation in fluctuating light environments. Nature communications, 2014. 5(1): p. 5439.

87. Davis, G.A., et al., Limitations to photosynthesis by proton motive force-induced photosystem II photodamage. elife, 2016. 5: p. e16921.

88. Karlsson, P.M., et al., The Arabidopsis thylakoid transporter PHT4;1 influences phosphate availability for ATP synthesis and plant growth. The Plant Journal, 2015. 84(1): p. 99–110.

89. Herdean, A., et al., A voltage-dependent chloride channel fine-tunes photosynthesis in plants. Nature Communications, 2016. 7(1): p. 11654.

90. Gabrys, H., et al., Mutants of Chloroplast Coupling Factor Reduction in Arabidopsis. Plant Physiology, 1994. 104(2): p. 769–776.

91. Lavaud, J., B. Rousseau, and A.-L. Etienne, In diatoms, a transthylakoid proton gradient alone is not sufficient to induce a non-photochemical fluorescence quenching. FEBS Letters, 2002. 523(1-3): p. 163–166.

92. Bailleul, B., et al., Electrochromism: a useful probe to study algal photosynthesis. Photosynthesis Research, 2010. 106(1): p. 179–189.

93. Pawlak, K., et al., On the PsbS-induced quenching in the plant major light-harvesting complex LHCII studied in proteoliposomes. Photosynthesis Research, 2020. 144(2): p. 195–208.

94. Li, X.-P., et al., Regulation of Photosynthetic Light Harvesting Involves Intrathylakoid Lumen pH Sensing by the PsbS Protein *. Journal of Biological Chemistry, 2004. 279(22): p. 22866–22874.

95. Kaminskaya, O.P. and V.A. Shuvalov, Biphasic reduction of cytochrome b559 by plastoquinol in photosystem II membrane fragments: evidence for two types of cytochrome b559/plastoquinone redox equilibria. Biochimica et Biophysica Acta (BBA)-Bioenergetics, 2013. 1827(4): p. 471–483.

96. Cardona, T., et al., Charge separation in Photosystem II: A comparative and evolutionary overview. Biochimica et Biophysica Acta (BBA) - Bioenergetics, 2012. 1817(1): p. 26–43.

97. Müh, F., et al., Light-induced quinone reduction in photosystem II. Biochimica et Biophysica Acta (BBA) - Bioenergetics, 2012. 1817(1): p. 44–65.

98. Satoh, K., et al., Binding affinities of benzoquinones to the QB site of Photosystem II in Synechococcus oxygen-evolving preparation. Biochimica et Biophysica Acta (BBA) - Bioenergetics, 1992. 1102(1): p. 45–52.

99. Moore, V. and W. Vermaas, Functional consequences of modification of the photosystem I/photosystem II ratio in the cyanobacterium Synechocystis sp. PCC 6803. Journal of Bacteriology, 2024. 206(5): p. e00454–23.

100. Kramer, D.M. and J.R. Evans, The importance of energy balance in improving photosynthetic productivity. Plant physiology, 2011. 155(1): p. 70–78.

101. Vass, I., et al., Reversible and irreversible intermediates during photoinhibition of photosystem II: stable reduced QA species promote chlorophyll triplet formation. Proceedings of the National Academy of Sciences, 1992. 89(4): p. 1408–1412.

102. Touloupakis, E., B. Cicchi, and G. Torzillo, A bioenergetic assessment of photosynthetic growth of Synechocystis sp. PCC 6803 in continuous cultures. Biotechnology for Biofuels, 2015. 8(1): p. 133.

103. Parry, M.A.J., et al., Rubisco activity and regulation as targets for crop improvement. Journal of Experimental Botany, 2012. 64(3): p. 717–730.

