## Supplemental Figures for "Adaptation of the *Chlorella* photosynthetic electron transport chain to environmental light conditions"

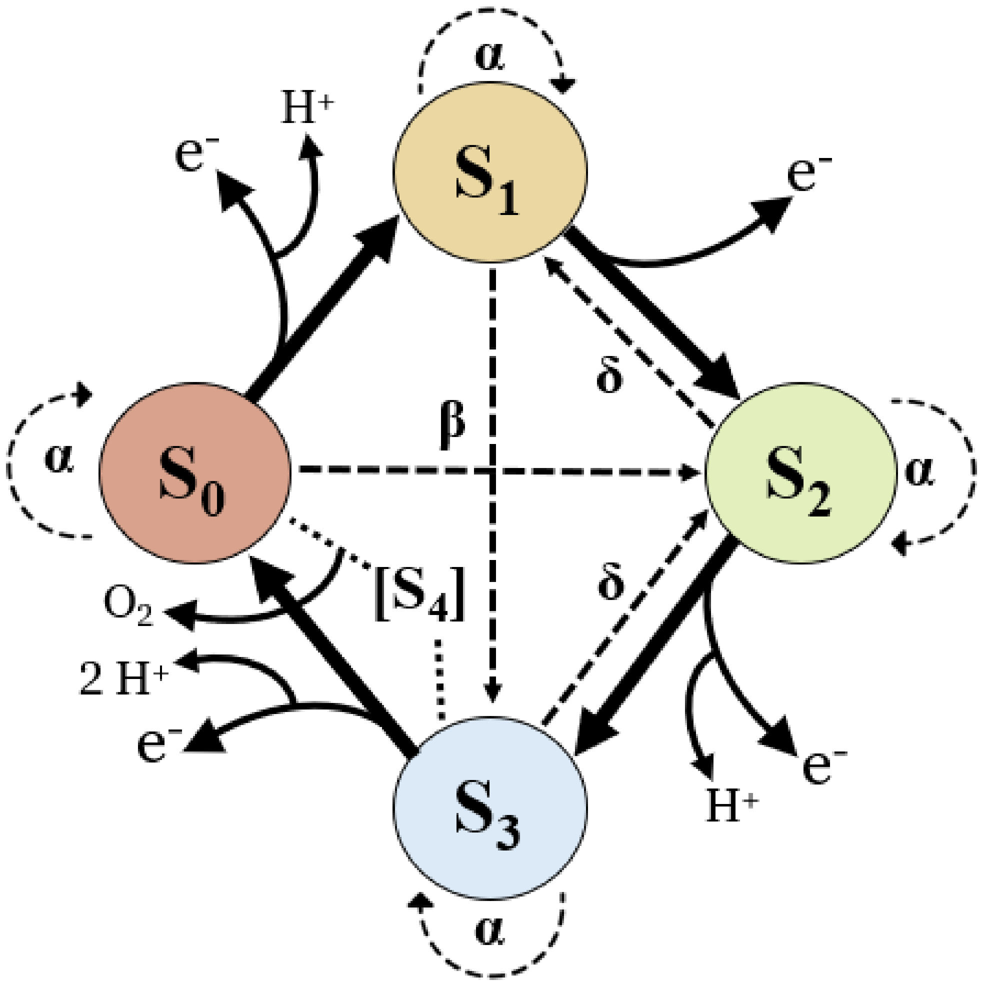


**Supplementary Figure S1**. A representation of the period-four oscillation of the WOC cycle, the byproducts of its progression, and its attributed inefficiencies modeled by the VZAD model. Each S-state is progressively oxidized, assuming successful charge separation of the reaction center.

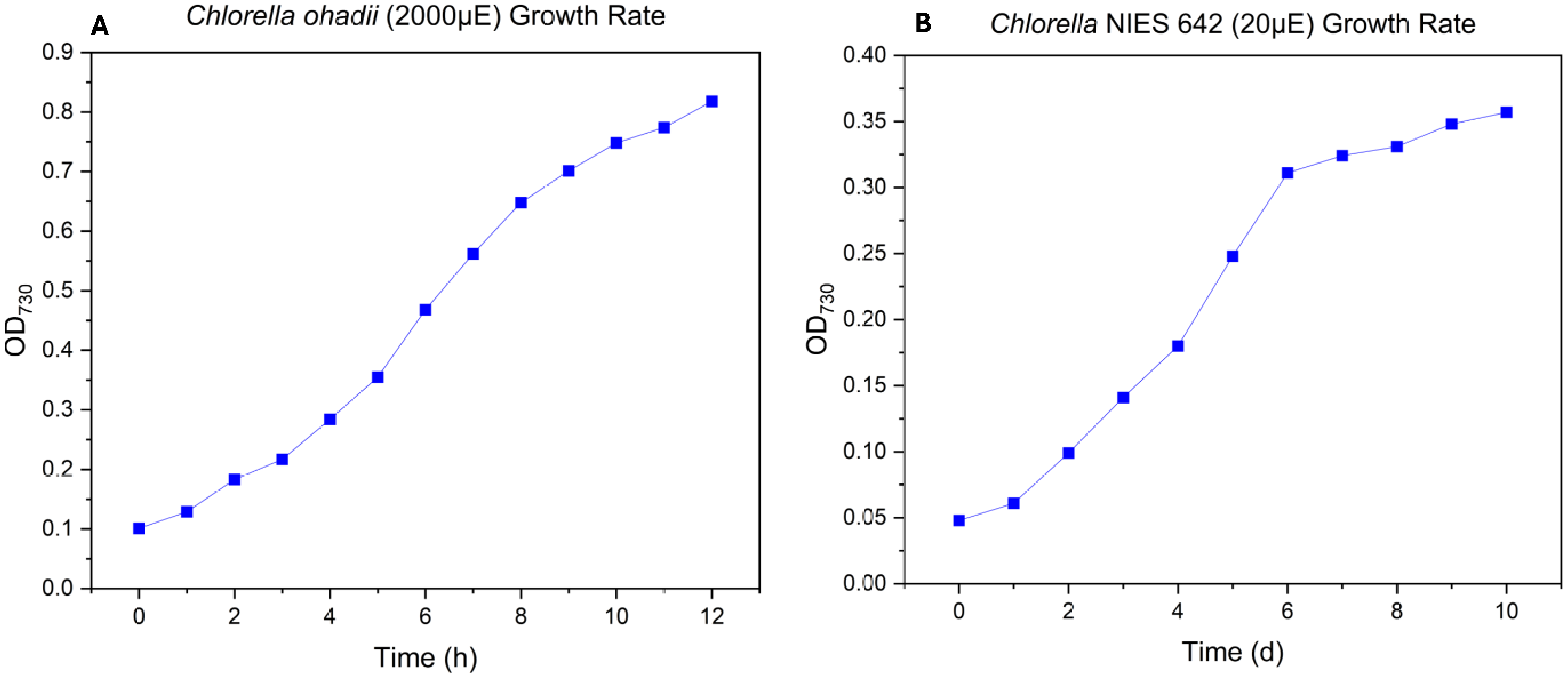


**Supplementary Figure S2**. Growth curves measured by optical density at 730 nm ($OD_{730}$) for two *Chlorella* species. Reported $OD_{730}$ values are an average of biological triplicate. **a)** *Chlorella ohadii*, grown in BG-11 medium at 30°C, bubbled with 1.5% $CO_2$,and a light intensity of 2000 µEin/m/s. **b)** *Chlorella* sp. NIES 642, grown in BG-11 medium at 30°C and a light intensity of 20 µEin/m/s.

**A**

| | 2,000 µEin | 20 µEin |
|---|---|---|
| | ***C. ohadii*** | ***C.* NIES 642** |
| Respiration | -69.3 | -11.8 |
| Oxygen Production | 1185 | 29.6 |
| Total $O_2$ Evolution | 1254 | 41.4 |
| | **µmol $O_2$/mg chl *a*/h** | |

**B**

| | ***C. ohadii*** | ***C.* NIES 642** |
|---|---|---|
| [chl *a*] (µg/mL) | 2.86 ± 0.11 | 9.14 ± 0.12 |
| [chl b] (µg/mL) | 0.43 ± 0.03 | 2.67 ± 0.05 |
| chl *a*:b ratio | 6.65 ± 0.53 | 3.42 ± 0.08 |

**Supplementary Figure S3**. Rate oximetry data for *Chlorella* species at their natural light intensities **a)** Respiration, $O_2$ production, and $O_2$ evolution (difference) for both species, normalized by chlorophyll *a* concentrations. **b)** Concentrations of chlorophyll (chl) *a* and b, and corresponding chl *a*:b ratio by species, extracted with chilled methanol as solvent.

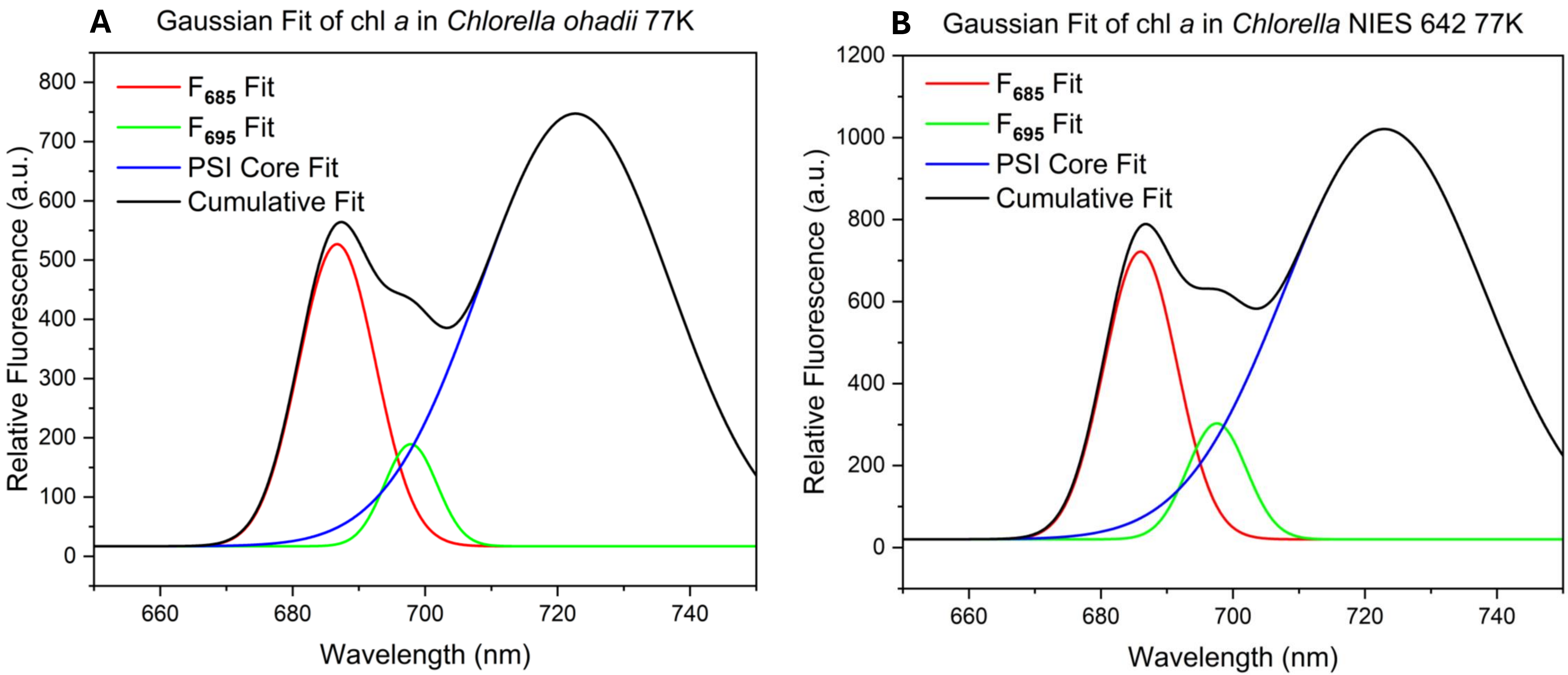


**Supplementary Figure S4**. Graphical representation of the Gaussian curves of photosystem components detailed in **Table 3**. Components fit are: $F_{685}$-active chlorophyll centers of PSII; $F_{695}$-chlorophyll trap usage (CP47); photosystem I.
